# Patient-derived organoid xenografts reveal the multifaceted role of the lncRNA *MALAT1* in breast cancer progression

**DOI:** 10.64898/2026.04.02.716096

**Authors:** Disha Aggarwal, Suzanne Russo, Kendall Anderson, Taylor Floyd, Raditya Utama, James A. Rouse, Payal Naik, Sara Pawlak, Shruti V. Iyer, Melissa Kramer, Shuchismita Satpathy, John E. Wilkinson, Qing Gao, Sonam Bhatia, Gayatri Arun, Martin Akerman, W. Richard McCombie, Alexey Revenko, Karen Kostroff, David L. Spector

## Abstract

Long non-coding RNAs (lncRNAs) have emerged as key regulators of tumor biology, however, thus far none have translated to cancer therapies. The lncRNA *MALAT1* is overexpressed in more than 20 cancers, including breast cancer and has been shown to function via various mechanisms in a context-dependent manner, in 2D cell lines and mouse models. However, its functional role and therapeutic potential have not been evaluated in clinically relevant patient-derived models. We investigated the therapeutic potential of *MALAT1-*targeting antisense oligonucleotides (ASOs) for breast cancer, using clinically relevant 3D human patient-derived organoids (PDOs) and PDO-xenograft (PDO-X) models. We systematically evaluated the efficiency of *MALAT1*-targeting ASOs using a biobank of 28 PDO models. Across three independent PDO-X models of triple negative breast cancer (TNBC), *MALAT1* depletion reproducibly drove widespread alternative splicing changes across all event types, with an enrichment of intron retention. Differentially spliced transcripts were enriched for targets of shared cancer-associated transcription factors, and *MALAT1* knockdown specifically altered the relative abundance of previously unannotated splicing isoforms. Beyond tumor-intrinsic effects, tumor-specific *MALAT1* depletion induced a consistent reduction in macrophage-associated gene signatures and reduced lung metastatic burden. Our data defines *MALAT1*’s multifaceted role in TNBC, coordinating alternative splicing, tumor-stroma crosstalk, and metastatic progression. Our study provides strong preclinical evidence supporting *MALAT1*-targeted ASO therapy and establishes PDO-X models as a clinically relevant platform for functional interrogation of TNBC therapies.

**Significance:** Using clinically relevant human PDO-X models, we show that *MALAT1* influences alternative splicing, metastatic potential and the tumor microenvironment. These data support the potential of oncogenic lncRNA *MALAT1* as a therapeutic target.

## Introduction

Breast cancer is the most common malignancy among woman, with approximately 2.3 million new cases reported globally in 2022 [1]. At current incidence rates, projections by the WHO Global Breast Cancer Initiative and the Lancet Breast Cancer Commission estimate that the incidence of new breast cancer cases will exceed 3 million cases annually by 2040 [2,3]. In the United States alone, Siegel et al. estimated 321,910 new breast cancer cases in 2026, accounting for ∼32% of all newly diagnosed cancers among woman [4]. Despite recent advances in early detection through annual screenings, targeted therapies, and improved understanding of genetic predisposition, breast cancer remains the second leading cause of cancer-related deaths among woman in the United States, with ∼42,140 deaths annually [4]. Breast cancer is broadly classified as ductal or lobular based on histopathological origin, and further classified based on the molecular expression of three clinically actionable receptors – hormone receptors i.e. estrogen (ER), progesterone (PR), and the human epidermal growth factor receptor 2 (Her2). Approximately 10-15% of breast cancers lack expression of all three receptors on the tumor cells. This triple-negative breast cancer (TNBC) subtype is among the most aggressive subtype of breast cancer and the lack of targetable ER, PR and Her2 receptors significantly limits available treatment options [5].

While protein-coding genes have entirely dominated the landscape of targeted therapeutics for breast cancer, non-coding RNA targets remain very much underexplored. According to the latest GENCODE annotation, 46% of annotated human genes encode long non-coding RNAs (lncRNAs), whereas only 25% of annotated human genes encode for all known proteins [6]. LncRNAs are generally defined as transcripts that are >200 nucleotides in length and lack open-reading frames required for translation to proteins/peptides [7]. Accumulating evidence indicates that lncRNAs regulate diverse cellular processes, including genome organization, gene expression, alternative splicing, epigenetic modification and mRNA stability [7]. There is a growing body of evidence supporting their roles in tumorigenesis, cancer progression and therapeutic resistance [8,9]. Among these, the lncRNA *Metastasis-Associated Lung Adenocarcinoma Transcript 1* (*MALAT1*) has been extensively studied and was one of the first cancer-associated lncRNAs identified [10–12]. Although our understanding of the roles of lncRNAs in cancer progression is rapidly expanding, these discoveries have not yet translated into clinical oncology, and thus far no lncRNA targets have advanced into clinical trials for any cancer type.

The distinct localization of *MALAT1* within nuclear speckles has long suggested a role in RNA processing [13]. Using 2D cell line models of development or cancer and mouse models of cancer, *MALAT1* has been shown to impact gene expression, alternative splicing and epigenetic regulation in a highly context-dependent manner [14–17]. *MALAT1* has been reported to directly associate with gene bodies and transcription termination sites of actively transcribing genes [18] and to interact with pre-mRNAs through associated proteins [19]. In specific cellular contexts, *Malat1* has been shown to interact with various nuclear proteins including transcription factors to modulate gene expression [20–23], as well as with transcriptional coactivators and histone modifiers to activate the cell growth gene expression program upon receiving mitogenic signals [22]. *MALAT1* has been implicated as an oncogenic lncRNA in >20 cancer types, including breast cancer [14,24,25], and has been shown to be involved in processes impacting various hallmarks of cancer [11,12,26,27]. In contrast, three studies contradicting a large body of evidence have proposed a tumor-suppressive role for *MALAT1* in breast cancer [28–30], highlighting unresolved context-dependent functions and underscoring the need to directly assess *MALAT1* function in clinically relevant model systems.

Previous work from our laboratory demonstrated *Malat1*-dependent changes in tumor transcription, alternative splicing, primary tumor differentiation and lung metastases in the MMTV-PyMT mouse model of breast cancer [14]. More recent studies have revealed an immunomodulatory role for *Malat1* in mammary tumor progression. Kumar et al. showed that deleting *Malat1* in the 4T1 syngeneic TNBC mouse model resulted in increased recognition and elimination of incipient metastatic cells by cytotoxic T-cells [24]. Additionally, a prior study showed that ASO-mediated *Malat1* knockdown in two syngeneic TNBC mouse models led to enhanced cytotoxic T-cell infiltration, and reduced macrophage infiltration of the mammary tumors and an augmented response to immune checkpoint blockade [25]. While mouse models of breast cancer highlight the potential of *Malat1* as an attractive target for breast cancer treatment, its function has not been examined in patient-derived xenograft (PDX) models. Given the complexity of breast cancer and the wide inter-patient diversity within TNBC [31–35], studying its relevance in patient-derived models is extremely important. Interestingly, a recent retrospective case study examining patient tumor sections over a breast cancer patient’s course of treatment and disease progression showed that *MALAT1* levels were consistently elevated in TNBC tumor cells relative to adjacent stroma, transiently reduced upon standard-of-care therapeutic interventions, and markedly increased in distant metastatic lesions [36].

In the present study, we investigated *MALAT1* function using clinically relevant human patient-derived models of TNBC. TNBC is well established as a highly heterogeneous disease, with variability observed across histopathological [31], transcriptomic [31–34,37–39] and genomic levels [31,35,40–43]. Patient-derived organoids (PDOs) are multicellular, three-dimensional (3D) structures derived from freshly resected patient tumors [44,45]. Breast tumor PDOs represent patient diversity and recapitulate the cellular heterogeneity and genetic features of the patient tumor, making them powerful pre-clinical models [46–48]. Our laboratory previously generated and characterized a biobank of TNBC PDOs and demonstrated engraftment to create PDO-derived xenografts (PDO-Xs) [46]. Here, we systematically screened PDOs from this biobank for *MALAT1* knockdown efficiency of human-specific *MALAT1*-targeting ASOs and established optimized PDO-X models for *in vivo* analysis. Using three independent TNBC PDO-X models, we demonstrate that ASO-mediated *MALAT1* knockdown induces patient-specific transcriptional responses. More importantly, perturbing *MALAT1* led to consistent disruption of alternative pre-mRNA splicing, with a pronounced enrichment of intron-retention events that generate novel transcripts. Interestingly, genes undergoing *MALAT1*-dependent splicing changes were significantly associated with a subset of transcription factors known to regulate gene expression in cancer. In addition, *MALAT1* depletion resulted in consistently reduced macrophage infiltration across all PDO-X models and significantly reduced lung metastatic burden. Together, these findings establish *MALAT1* as a multifaceted regulator of human breast cancer progression and provide strong support using clinically relevant patient-derived model systems, for pursuing *MALAT1* as a therapeutic target. The present study is a first of its kind in successfully utilizing patient-derived breast tumor organoids (PDOs) and PDO-xenograft models for a comprehensive mechanistic perturbation study of a potential therapeutic target.

## Materials and Methods

### Patient Material

Tumors from breast cancer patients as well as normal breast tissue from individuals undergoing reductive mammoplasty were obtained from Northwell Health to generate organoids as previously published [46]. Samples were acquired in accordance with Institutional Review Board protocol IRB-03-012 and IRB 20-0150 with written informed consent from the patients. Formalin-fixed paraffin-embedded (FFPE) slides of patient tumor tissues were also obtained from Northwell Health Biorepository. Specific information including findings in the anonymous patient pathology reports for all samples is available in Table S1. The collection of genomic and phenotypic data was consistent with HHS 45 CFR Part 46 (Protection of Human Subjects) and the NIH Genomic Data Sharing (GDS) Policy.

### Organoid culture

A biobank of breast tumor organoids was previously established and expanded in culture in our laboratory [46] using a protocol [49] that was adapted from the Clever’s lab protocol [47]. Organoid models labeled with the prefix HCM-CSHL were developed as part of the Human Cancer Model Initiative (HCMI; https://ocg.cancer.gov/programs/HCMI) and a subset of those models are or will be available for access from ATCC.

### *In vitro* ASO treatment

Specific 16-mer 3-10-3 gapmer antisense oligonucleotides (ASOs) with a phosphorothioate backbone and constrained ethyl (cEt) modified nucleotides on the 5’ and 3’ ends were designed by Ionis Pharmaceuticals, Inc for targeting human *MALAT1* RNA. The control Scrambled ASO was also of the same chemistry designed by Ionis Pharmaceuticals to have no significant sequence homology to known human genes (Table S3). The PDOs were plated in a 100 µl dome of Matrigel (Corning 356231) in 12 well plates (VWR 82050-926). Once the domes solidified, the culture medium supplemented with 3 µM of *MALAT1*-targeting ASO was added to the organoids. Untreated organoids with medium (labeled mock) served as the control. An additional ASO control used medium supplemented with 3 µM Scrambled ASO (ScASO). The medium and ASOs were replenished on day 3 and organoids were harvested to assess *MALAT1* knockdown efficiency on day 6 by adding 1 ml TRIzol reagent (Life Technologies 15596018) per well containing organoids in the Matrigel dome for extracting RNA.

### Organoid Histology

Matrigel domes containing organoids were scraped and transferred to a 1% Bovine Serum Albumin (BSA) (Gibco 15260037) pre-coated 50 ml conical tube. The Matrigel-embedded organoids were centrifuged at 400 x g for 5 minutes and washed with cold 1X DPBS. Organoids were harvested using cold Organoid Harvesting Solution (Trevigen 3700-100-01) at a ratio of 5x the initial volume of Matrigel, at 4°C for 30 minutes on a shaker and the pellet of organoids was washed and fixed using freshly prepared 4% formaldehyde (Thermo Scientific 28908) at room temperature for 30 minutes. After two washes, the organoids were resuspended in 1 ml 1X DPBS and transferred to a BSA coated 1.5 ml Eppendorf tube. After spinning, the pellet was embedded in 2% agarose. The agarose mold containing organoids was paraffin-embedded and 5 µm serial sections were cut to be used for H&E or IHC or smRNA-FISH.

### Animals

All animal procedures and studies were carried out in accordance with the CSHL Animal Care and Use Committee (IACUC Protocol No. 2021-1197). Six-week-old female NOD *scid* gamma (NOD.Cg-*Prkdc^scid^ Il2rg^tm1Wjl^*/SzJ) mice were purchased from the Jackson Laboratory (005557). All organoid injections were done after acclimating the mice at the Cold Spring Harbor Laboratory’s Animal Shared Resource facility for a week.

### PDO-X generation and *in vivo* ASO treatment

Confluent wells of organoids were harvested using TryPLE (Life Technologies 12605028). After trypsinization, the organoids were resuspended in 1x DPBS and filtered using a 70 µm filter (Pluriselect 43-50070-51) and counted using Trypan blue (Gibco 15250061) and the cell countess (Countess II FL, Life Technologies). The organoids were resuspended in 1:1 mix of Matrigel and sterile 1X DPBS at a concentration of 2-4 million cells/100 µl based on the growth rate of the organoid line to be injected. The organoids were placed on ice. The mice were anesthetized with 1.5-2% isoflurane, weighed and 2-4 million cells i.e. 100 µl injected into the mammary fat-pads #4 (both left and right) of 6-week-old NOD *scid* gamma (NOD.Cg-*Prkdc^scid^ Il2rg^tm1Wjl^*/SzJ) mice.

Mice were monitored and primary tumors were measured every week using vernier calipers. Once the tumors were palpable, the mice were anaesthetized and treated with either a Scrambled control ASO or one of two *MALAT1*-targeting ASOs subcutaneously at 50 mg/kg/day twice a week. The blinded ASO treatment continued until the endpoint. The animals were sacrificed when at least one of the two tumors reached 2 cm in size as measured externally by Vernier Calipers. The tumor volume was measured as (length x (width)^2^)/2 with width being the smaller measurement. The tumors (right and left), lungs, brain, liver, spleen, kidneys and lymph nodes were collected from each animal into labeled cassettes during necropsy. Each lobe of the lung was separated while placing it in the cassette. The tumors were cut into two pieces. One part was snap frozen on dry ice and stored at –80 °C thereafter for RNA extraction. The other half was fixed in formaldehyde along with other tissues overnight in freshly prepared 4% formaldehyde (Thermo Scientific 28908) after which they were washed twice with 1X DPBS. They were stored in 1X DPBS at 4 °C until embedding. Fixed tissues were processed and embedded in paraffin blocks by the histology core facility at Cold Spring Harbor Laboratory.

### Hematoxylin and eosin (H&E) staining and immunohistochemistry (IHC)

Hematoxylin and eosin (H&E) staining and immunohistochemistry (IHC) were performed at the CSHL Histology Core Facility. Tissue samples fixed in 4% formaldehyde were processed using a Thermo Excelsior ES tissue processor and embedded with a Thermo HistoStar embedding system, following the manufacturer’s protocols. Paraffin blocks were sectioned at 5 µm and mounted onto positively charged slides (Fisherbrand™ Superfrost™ Plus Microscope Slides). For H&E staining, slides were processed on a Leica Multistainer (ST5020). Briefly, slides were deparaffinized, rehydrated, and stained with hematoxylin (Hematoxylin 560 MX, Leica) for 1 minute. This was followed by destaining in Define MX-aq (Leica) for 30 seconds, bluing in Blue Buffer 8 (Leica) for 1 minute, and counterstaining with eosin (EOSIN 515 LT, Leica) for 30 seconds. After dehydration, coverslips were applied using a Leica CV5030 robotic coverslipper.

For IHC staining, a Roche Discovery Ultra automated stainer was used following standard protocols. Antigen retrieval was performed using Benchmark Ultra CC1 (Roche) at 96°C for 1 hour. Primary antibodies were incubated at 37°C for 1 hour, and immunosignals were detected and amplified using the Discovery OmniMap HRP detection system (Discovery DAB, Roche). The following antibodies were used: anti-human phosphorothioate (anti-ASO) antibody (Ionis Pharmaceuticals in-house, dilution 1:3,000), anti-human mitochondria antibody (Millipore MAB1273, 1:400), F4/80 antibody (Cell Signaling Technology, #70076, 1:500), anti-ER antibody (Abcam, ab27595, 1:2). The IHC data analysis was done using QuPath software quantifying the DAB stained area (brown) against a hematoxylin background stain.

### RNA isolation

The tissues were homogenized to a powder while still frozen using the Qiagen TissueLyzer II machine by shaking at 30 rpm for 60 seconds. Total RNA was extracted from powdered tissue or organoids in Matrigel using TRIzol (Life Technologies 15596018) following the manufacturer’s protocol.

### Reverse transcription/cDNA synthesis and quantitative real-time polymerase chain reaction (qRT-PCR)

For reverse transcription, 1 µg of total RNA was used for cDNA generation post DNase I treatment using random hexamer primers and the TaqMan reverse transcription reagent kit (ThermoFisher A25743). For qRT-PCR, 1:10 diluted cDNA was mixed with 2X SyBr Green (ThermoFisher A25743) and target-specific primers. Details of sequences for the qRT-PCR primers used in this experiment are listed in Table S4.

### Library construction and RNA sequencing

The quality and integrity of the extracted RNA was assessed via Agilent Tapestation 4150. All samples with a threshold of ≥8 RNA integrity number (RIN) were submitted for library preparation and sequencing. Stranded, poly(A)+ RNA libraries were prepared using the KAPA mRNA HyperPrep kit (Roche 08098123702) at the Cold Spring Harbor Laboratory NextGen Sequencing core facility and sequenced paired-end 150 bp on NextSeq 2000 P4 or Singular G4 F3 platforms (NH048T PDO-Xs). Other sets of samples (DS115T and NH85TSc PDO-Xs) were processed for library preparation at Novogene using the NEBNext Ultra II directional library prep kit for Illumina (E7760L) and sequenced on Novaseq X Plus 25B platform for an output of 50 million reads per sample. Each set had internal controls processed together.

### RNA sequencing differential gene expression analysis

Bulk RNA-seq analyses were implemented and integrated using the CodeSpringLab platform developed by the CSHL Bioinformatics core facility, downloaded from https://github.com/RadUtama/CodeSpringLab.git. Quality control of raw sequencing files (fastq) was implemented using FastQC (v0.11.8) and FastQ Screen (v0.15.2) [50] with default parameters. Sequence alignment was performed using STAR (v2.7.10a) [51] with parameters set to --outFilterMismatchNmax 2 –outFilterMultimapNmax 2 --outSAMtype BAM SortedByCoordinate --outSAMunmapped None --outSAMstrandField None. The human genome reference (FASTA assembly and GTF annotation) was extracted from GENCODE GRCh38.p14 (release v47) and the mouse genome reference (FASTA assembly and GTF annotation) was extracted from GENCODE GRCm39 (release M36). The reads were combined into a combined reference genome used for alignment of all PDO-X RNA-seq experimental data.

Strandness was estimated using RSeQC infer_experiment.py (v4.0.0) [52]. Gene quantification was performed using featureCounts from Subread (v2.0.2) [53] with parameters set to -p –countReadPairs -t exon -Q 12 -C –minOverlap 1. Gene normalized counts in TPM were calculated using RSEM (v.1.3.3) [54]. Differential analysis was performed using DESeq2 (v1.36) [55] and log-fold shrinkage apeglm package [56], with parameters set to a minimum of 10 read counts for each gene summed over all samples. Differentially expressed genes (DEGs) were defined as genes with p-value< 0.01.

The DEGs common (in the same direction i.e. up or down) between the two different *MALAT1*-targeting ASOs were identified using the dplyr [57] package in R [58]. These common gene lists of up or downregulated genes were used separately as input in Enrichr [59] to identify the pathways and cell types (p-value<0.05) that are enriched among the differentially expressed genes.

### Splicing data analysis

RNA-sequencing (RNA-seq) data were processed through FastQC to evaluate sample quality, remove adapter sequences and remove short (<40 nucleotides) and poor-quality reads. Filtered reads were aligned to either the GENCODE version 43 (GRCh38.p13) or RefSeq Release 218 (GRCh38.p14) human genomes using STAR.

RNA-seq sample quality was assessed using an internal quality control assessment tool that leverages quality control metrics generated by FastQC and STAR. The samples were then analyzed using the SpliceCore^®^ commercial software platform to produce alternative splicing (AS) profiles between each *MALAT1*-targeting ASO vs Scrambled ASO datasets. The SpliceCore platform uses the SpliceTrap™ algorithm [60] to align RNA-seq data to a database of known splicing events (TXdb) and quantify the “percent spliced in” (PSI) values for each AS event. Next, SpliceDuo™ was utilized for case/control comparisons and reporting splicing changes as ΔPSI values ranging from - 1 (complete exon skipping) to 1 (complete exon inclusion) [61]. Significant AS changes identified by SpliceDuo (p-value <0.1) were filtered (|dPSI| ≥ 0.2, % reproducible samples ≥ 33%, consistency > 90%) to identify biologically relevant differences in splicing between cases and controls in *in vivo* organoid experiments and splicing event type was assigned based on the AS change - alternative acceptor (AA), alternative donor (AD), cassette exon (CA) or intron retention (IR). The distribution of splicing event types (AA, AD, CA, IR) for the filtered splicing events was compared between each organoid sample and TXdb-curated (known) splicing events. P-values were generated using the Chi-Squared Goodness of Fit test.

Genes mapping to splicing events that passed filters were submitted to the Enrichr webserver database, and significantly (p-value<0.05) enriched transcription factors (TFs) associated with alternatively spliced transcripts for each PDO-X experiment were identified using the ChEA 2022 database. The overlap between significantly-enriched TFs for each ASO for individual PDO-X models was first determined and then the overlap between the three independent PDO-X experiments was shortlisted. The identified list of 15 TFs was input into Enrichr to identify pathways and diseases they regulate via KEGG 2026 and Jensen Diseases Curated 2025 databases respectively (p-value<0.05).

### Affinity-based Cas9-Mediated Enrichment (ACME) and long read sequencing

High Molecular Weight (HMW) DNA was extracted from 50-100 µl of thawed organoid pellets using the ‘Extraction from Cells’ Monarch kit (NEB Catalog #T3050) and quantified using the Qubit fluorometer (Thermo Fisher Scientific) and integrity was assayed using Femto Pulse System (Agilent). Four to six crRNA guides (Table S5) were designed per target using the CHOPCHOP webtool (http://chopchop.cbu.uib.no/) [62]. Using 10 µM pooled guide RNAs, ACME was performed using the Oxford Nanopore Technologies (ONT) kit (CS9109) and the previously reported ACME modification protocol was incorporated after the Cas9 cleavage step [63]. Libraries were prepared for loading as per the manufacturer’s protocol and ∼ 50 fmol of the prepared library was loaded on FLO-MIN106 R9.4.1 flowcells with >1200 active pores. Platform QC was performed on the GridION sequencer. SQK-CS9109 kit was chosen for live base calling using the high accuracy model. For the NH048T PDO and blood samples, a modified version of ACME [63] was adapted to be compatible with the newer R10 ONT chemistry (Table S6). These two samples were sequenced on the PromethION high throughput instrument using FLO-PRO114M R10.4.1 flowcells, with >6000 active pores following Platform QC. SQK-LSK114 kit was chosen on the MinKNOW interface.

### Long read sequencing analysis

ONT reads for each organoid sample were aligned to the hg38 reference genome using minimap2 [64]. Structural variants (SVs ≥ 30 bp) were called using Sniffles2 [65], collapsed for events within 1 kb of each other and filtered for ≥25% read depth. SVs were annotated using UCSC genome browser tracks, including dbVar/DGV, ENCODE regulatory elements, OMIM genes, eQTL markers, and segmental duplications.

Single-nucleotide variants (SNVs) were called using Clair3 [66] with default parameters and filtered for a minimum phred scaled variant quality score of 20. SNVs were compared to matched previously generated Illumina short read sequencing of a targeted gene panel [46] using bedtools [67] to assess overlap. DNA samples from patient’s blood cells (suffix ‘Bl’) were used as matched germline reference to identify tumor-specific variants.

Base modifications were called using the ONT raw signal data (fast5/pod5 files). Briefly, Guppy or Dorado super accuracy base calling was run using the model to detect 5-methylcytosine (5mC) and 5-hydromethylcytosine (5hmC) in CpG context, and to align the reads to the hg38 reference. The resulting BAM files were then assessed using modbamtools [68] to visualize per-read CpG methylation patterns (red for methylated CpG and blue for unmethylated CpG) across loci of interest.

### Transmission Electron Microscopy

The tumor tissue was harvested and placed immediately into freshly prepared 2.5% glutaraldehyde and 2% formaldehyde in 0.1 M sodium phosphate buffer fixative. The samples were fixed overnight at 4 °C. The samples were washed with sodium phosphate buffer then post-fixed with 1% osmium tetroxide in 0.1 M phosphate buffer. Tissues were dehydrated in a graded series of ethanol (50%, 70%, 95%, 100% x2). The transitional solvent, propylene oxide was used, and samples were gradually infiltrated with EMbed 812 (Electron Microscopy Sciences) resin. Tissue samples were left in 100% resin overnight and changed to fresh resin for an hour before embedding in BEEM capsules. The capsules were polymerized overnight at 65 °C in a vacuum oven. The samples were trimmed and sectioned on a Reichert Ultracut E using a diamond knife. Thick sections were stained with 1% toluidine blue and areas of interest chosen under light microscopy. The chosen resin blocks were further trimmed, thin sectioned, and mounted on copper support grids. The prepared grids were post stained with 4% alcoholic uranyl acetate for 10 minutes followed by Sato’s triple lead stain for 2 minutes. The images were captured on a Hitachi HT7800 120kV transmission electron microscope operated at 80kV equipped with the AMT Nanosprint 15, 15-megapixel digital camera.

### Single-molecule RNA fluorescence *in situ* hybridization (smRNA-FISH)

smRNA-FISH was performed on 5 µm thick FFPE tissue sections of the patient tumors, obtained from Northwell Health Biorepository, using a previously described protocol as per manufacturer’s protocol using *MALAT1*-targeting probes [36].

### Statistical Analysis

Statistical analyses were performed using R on RStudio and Microsoft Excel. Specific tests are indicated in the figure legends along with the statistical significance outputs.

### Materials, data, and code availability

All raw RNA sequencing data will be available to download from dbGaP (phs004666.v1.p1).

## Results

### *MALAT1*-targeting ASOs have higher knockdown efficiency in fast-growing breast tumor organoids derived from high-grade, poorly differentiated tumors

We first optimized ASO-mediated knockdown of *MALAT1* in three-dimensional patient-derived breast tumor organoid (PDO) models. The ASOs used in this study are 16-nucleotide gapmers with a central DNA stretch of ten nucleotides flanked by three ribonucleotides on each end (Figure 1A). The sugar moieties have a constrained ethyl (cEt) modification and a phosphorothioate backbone. The gapmer chemistry is crucial for eliciting a RNase H1 activity on the DNA-RNA hybrid generated upon the ASO binding to its complementary RNA target. RNase H1 in combination with exonucleases degrades the target RNA [69] which in this case is *MALAT1* (Figure 1A).

**Figure 1:**
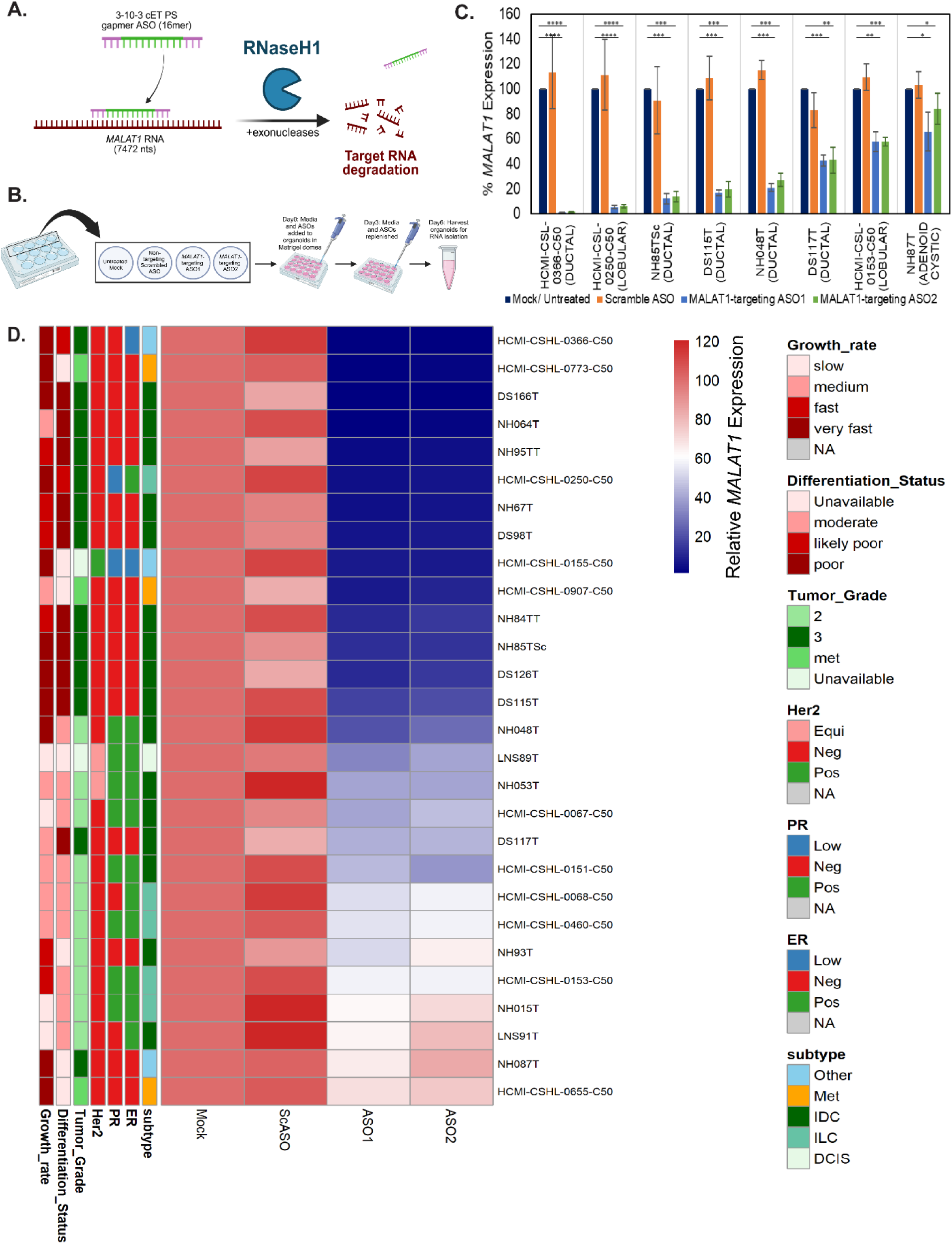
Screening patient-derived breast tumor organoid models (PDO) for antisense-oligonucleotide (ASO) mediated knockdown (KD) of *MALAT1*. (A) Schematic for ASO design and mechanism of action (Created in BioRender. Aggarwal, D. (2025) https://BioRender.com/eu45n3n). ASO Design: 16-nucleotides (16-mer) long gapmer ASO i.e. 3 ribonucleotides flanking the center 10 deoxyribonucleotides (3-10-3) with 2’-O-ethyl modifications (cEt) and a phosphorothioate (PS) backbone. (B) Schematic of the experimental design for ASO-mediated *in vitro MALAT1* KD. (C) *MALAT1* expression as measured by qRT-PCR. X-axis represents organoid lines with pathological subtype in parentheses. Error bars represent SEM. (p-value: ns-not significant, *<0.05, **<0.01, ***<0.001, ****<0.0001 as measured by Student’s t-test: two sample assuming unequal variances) (D) *MALAT1* expression in 28 PDO models as measured by qRT-PCR. Growth rate represents *in vitro* organoid growth in culture. Differentiation status of the tumor, tumor grade, Her2, PR, ER expression and breast cancer subtype information as per patient clinical pathology report. Neg – negative, Pos – positive, Equi – equivocal, met – axillary lymph node recurrence, ScASO – scrambled ASO (control), ASO1 and ASO2 – *MALAT1* targeting ASOs, ER – Estrogen Receptor, PR – Progesterone Receptor, Her2 – Human Epidermal Growth Factor Receptor 2, IDC – invasive ductal carcinoma, ILC – invasive lobular carcinoma, DCIS – ductal carcinoma *in situ*, Met – lymph-node metastasis-derived model, Other – rare subtype of breast cancer.

Two independent *MALAT1*-targeting ASOs were evaluated for their knockdown efficiency following “free uptake” (delivery of unformulated ASOs) from the culture medium by PDOs in Matrigel domes (Figure 1B). The initial eight PDO models tested presented with a variable efficiency of ASO-mediated *MALAT1* knockdown compared to untreated organoids (Figure 1C). A scrambled ASO was used as a non-targeting ASO control. Therefore, we expanded the number of models to evaluate *MALAT1* knockdown efficiency in 28 PDO models, and the results are summarized in Table S1. Of the 28 models, ∼54% (15/28) models exhibited a >70% *MALAT1* knockdown efficiency with either of the two ASOs at 3 µm ASO concentration. We observed that the PDO lines with >70% knockdown efficiency, typically exhibited rapid growth *in vitro* (Figure 1D, above black line). Examination of the corresponding patient pathology reports revealed a striking trend among the PDO models that display the strongest ASO-mediated *MALAT1* knockdown efficiency: these PDO models were most often derived from high grade, poorly differentiated, aggressive tumors, most of which are TNBC (Figure 1D).

To examine if endogenous *MALAT1* RNA level was a limiting factor impacting the ASO knockdown efficiency, we reanalyzed the bulk RNA-sequencing data previously generated to characterize the organoid models [46]. As seen in Figure 2A, low knockdown efficiency does not inversely correlate with high endogenous *MALAT1* expression in the PDO lines with <70% *MALAT1* KD. Further, using previously generated bulk RNA-seq data [46], we compared PDO models with high knockdown efficiency to PDOs with low knockdown efficiency. We observed 3,029 genes were differentially expressed (p-value<0.01) of which 1,714 genes are higher and 1,315 gene are lower in the PDO models with high *MALAT1* knockdown efficiency (Figure 2B). Pathway analysis via Enrichr (p-value<0.05) revealed that the genes enriched in the PDO models with strong *MALAT1* knockdown efficiency are significantly represented by cell cycle-associated pathways (Figure 2C and 2D). Using an antibody that targets the phosphorothioate backbone of ASOs (produced by Ionis Pharmaceuticals), we performed immunohistochemistry (IHC) on 5 µm sections from two *in vitro* cultured PDO models. The PDO lines chosen show variable *MALAT1* KD efficiency via qRT-PCR (Figure 1D). As is evident from Figure 2E, the ASOs enter the cells of both organoid lines, irrespective of the organoid size or downstream knockdown efficiency. Additionally, we also assessed whether endogenous RNase H1 expression differed between the PDO lines that gave strong *MALAT1* KD vs the lines with weak KD. We found that not all PDO lines with a strong *MALAT1* KD had a high RNase H1 transcript expression compared to the lines with <70% *MALAT1* KD efficiency (Figure S1A). Further, we also observed no difference in ASO knockdown efficiency based on the size of the organoids at the time of seeding when ASOs are first administered (Figure S1B-D).

**Figure 2:**
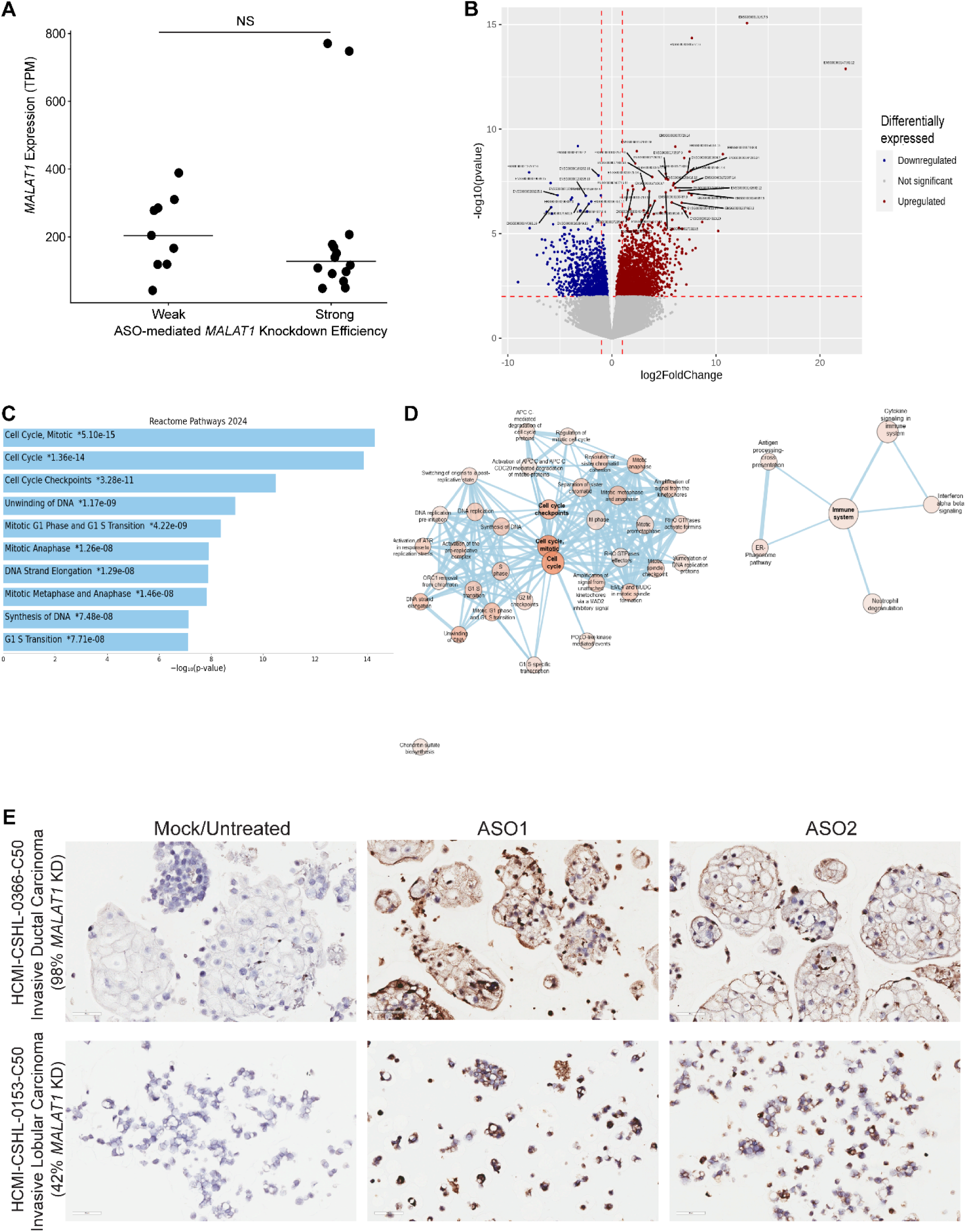
Assessing molecular factors and their correlation with *MALAT1* knockdown (KD) efficiency in breast tumor PDOs. (A) Endogenous *MALAT1* transcripts per million (TPM) for organoid lines with variable knockdown efficiency (NS – not significant as measured by Wilcoxon rank-sum test, p-value=0.2439). (B) Volcano plot showing differential gene expression comparing organoids with >70% *MALAT1* KD efficiency to PDOs with <70% *MALAT1* KD (p-value<0.01). (C) Pathway analysis using Reactome database via Enrichr (p-value<0.05) showing pathways enriched in PDO models with >70% *MALAT1* KD efficiency. (D) Network pathway analysis via Cytoscape, showing significantly enriched pathways associated with cell cycle. (E) Immunohistochemistry using an anti-ASO (phosphorothioate backbone) antibody to label 5 µm sections of PDOs. ASO1 and ASO2 – *MALAT1* targeting ASOs.

### TNBC patient-derived organoid xenografts (PDO-Xs) recapitulate the patient’s tumor

A previous study from our lab generated PDO-Xs from eight PDO models by injecting 50,000 organoids in the fat-pads of Nude immunocompromised mice [46]. This strategy resulted in limited success where three of the eight lines successfully grafted [46]. Here, we optimized this strategy to examine the take-rate of ten PDO models in NOD *scid* gamma (NSG) (NOD.Cg-*Prkdc^scid^ Il2rg^tm1Wjl^*/SzJ) mice. Based on the *in vitro* growth rate of the organoid line, 2-4 million cells from digested PDOs (mixture of single cells and small organoids) were bilaterally injected into mammary fat-pads #4 of NSG mice (Figure 3A). The success rate was observed to be much higher with this optimized strategy. All organoid models led to palpable tumors upon engraftment, with 83-100% take rate (Figure 3B). Additionally, we observed that different PDO lines have widely variable tumor growth rates *in vivo*. Some PDO models consistently form measurable tumors (∼100 mm^3^) within 2 weeks of transplantation, whereas some PDO models lead to palpable tumors after 15 weeks of transplantation. Additionally, tumor growth rates vary markedly between models: some tumors reach maximum growth of 2 cm within 13 weeks post-transplantation, whereas others do not achieve comparable sizes even after 31 weeks (Figure 3C).

**Figure 3:**
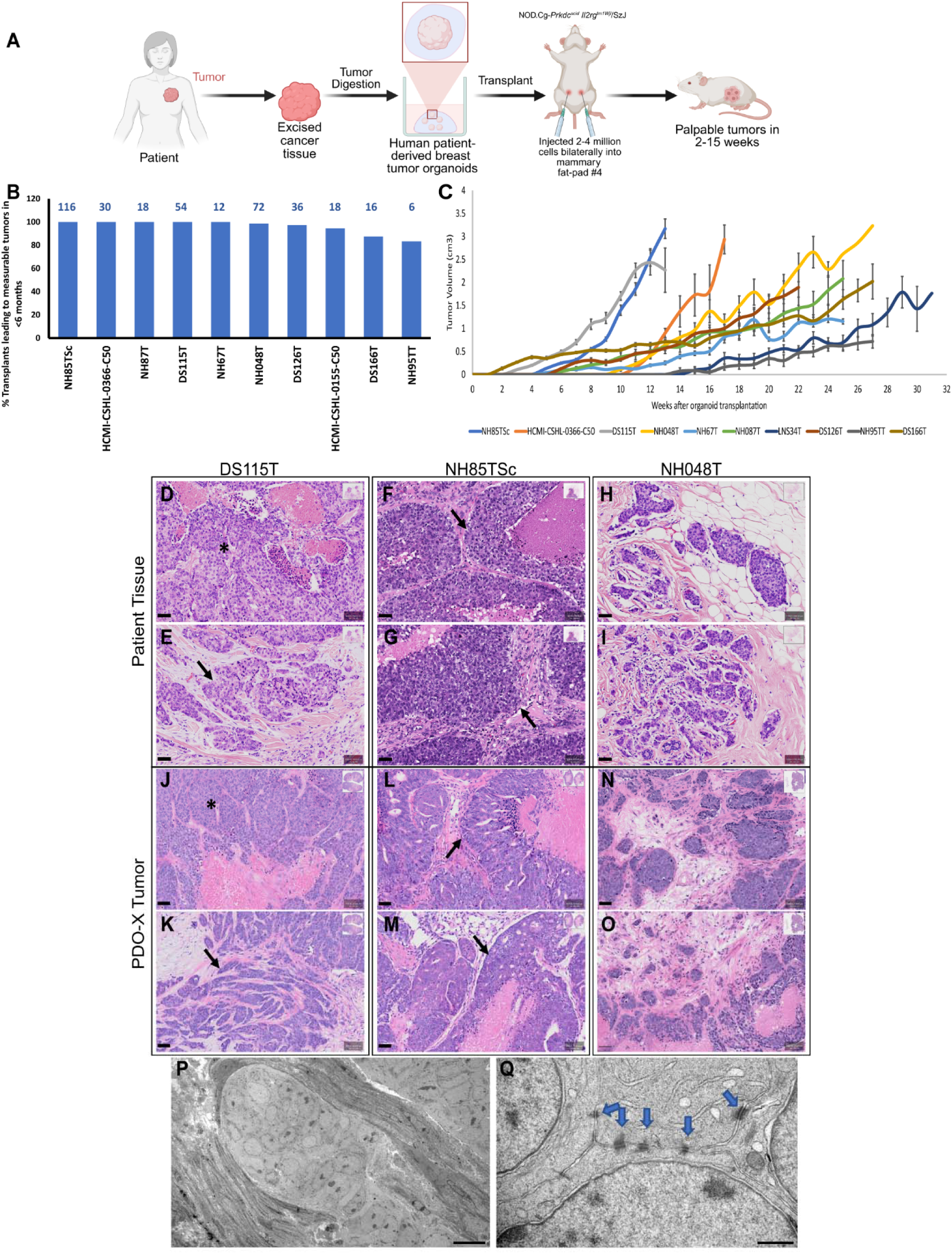
Generation of patient-derived organoid xenografts (PDO-Xs). **(A)** Schematic representation of the adopted methodology to generate breast cancer PDO-Xs (Created in https://BioRender.com). (B) Success rate of PDO transplantation into NOD *scid* gamma (NSG) mice. The numbers above the bars indicate number of transplants attempted. (C) Average palpable tumor volume of PDO-X tumors over time. (D-O) Hematoxylin and eosin (H&E) staining of original patient tumor sections (D-I) and corresponding PDO-X tumor sections (J-O) in NSG mice for DS115T, NH85TSc and NH048T PDO-X models. Scale bar = 50 µm. (P-Q) Transmission electron microscopy images of DS126T PDO-X tumors. (P) Scale bar = 10 µm. (Q) Scale bar = 600 nm. Blue arrows: cell-cell junctions between tumor cells.

Here, we compared the primary tumors from newly established PDO-X models in NSG mice (Figures 3J-O and S2G-L) to the original patient tumors (Figures 3D-I and S2A-F) from which the corresponding organoid models were derived. Hematoxylin and eosin (H&E) staining revealed that the PDO-X tumors recapitulate overall morphology of the original patient tumors (Figures 3D-O and S2A-L). DS115T patient tumor tissue (Figure 3D, E) and its corresponding PDO-X tumor (Figure 3J, K) contained solid and invasive areas. The patient tissue solid regions were composed of sheets of small cells with pale cytoplasm and the PDO-X tumor recapitulated this pattern but showed increased squamous differentiation. NH85TSc patient (Figure 3F, G) and PDO-X tumors (Figure 3L, M) were phenocopies with both displaying stratified tumor cells separated by bands of connective tissue and highly necrotic areas. NH048T patient (Figure 3H, I) and PDO-X tumors (Figure 3N, O) were each characterized by cords and nests of poorly differentiated cells infiltrating a rich connective matrix, with closely matching overall architecture.

DS126T patient (Figure S2A, B) and PDO-X tumor (Figure S2G, H) resembled a typical invasive carcinoma with poor differentiation and squamous features. DS166T patient tumor (Figure S2C, D) and the matched PDO-X tumor (Figure S2I, J) contained large areas of spindle cells interspersed with nests of poorly differentiated epithelial cells, reproducing the tumor’s biphasic morphology. HCMI-CSHL-0155-C50 patient (Figure S2E, F) and PDO-X tumor (Figure S2K, L) pairs showed similar morphology: the tumor displayed stratified tumor cells around necrotic centers within loose connective tissue, while the PDO-X tumor formed small cords of tumor cells with squamous differentiation, mirroring the patient tumor’s structural pattern. In summary, patient tumors and their PDO-X counterparts largely phenocopied the histopathological features, faithfully reproducing key architectural patterns and differentiation states with only minor variations in squamous features.

To further characterize the PDO-X tumors at higher resolution, we evaluated the PDO-X tumors from a fast-growing organoid line, DS126T, using transmission electron microscopy (TEM) (H&E - Figure S2G, H). The densely packed cells of the tumor were infiltrated by mouse stromal cells (Figure 3P). The interface of stroma and tumor indicated the presence of extracellular matrix fibers, vacuoles and lipid granules (Figure 3P). The stroma contained cells of different types (Figure 3P). At a higher magnification, we were also able to visualize extensive cell-cell junctions between the densely packed tumor cells (Figure 3Q, arrows).

### Transcriptomic and genomic characterization of PDO models

Bulk RNA-seq analysis comparing all previously characterized [46] PDO models compared to normal breast organoids derived from reduction mammoplasties, showed a strong clustering of all normal organoid models (red dots) away from tumor models (blue dots) (Figure 4A). The three PDO models selected for *in vivo* studies cluster apart from each other (circled), thus representing the wide patient/subtype diversity within TNBC (Figure 4A). Importantly, different passages of the same organoid line clustered together (circled, Figure 4A). Additionally, two PDO models generated from the same patient collected from primary tumor (NH85TSc) or later from lymph node recurrence (HCMI-CSHL-0907-C50) also clustered together (red circle, Figure 4A). Overall, this highlights the previously identified wide patient-to-patient variability in breast cancer [31–34,37–39], and the transcriptional consistency of specific tumor models. The three PDO-X models – NH85TSc, DS115T and NH048T were some of the fastest growing models *in vivo* (Figure 3C). NH85TSc and DS115T patient tumors were histologically classified as TNBC invasive ductal carcinoma (IDC) whereas, the NH048T patient tumor was ER+PR+Her2. However, the organoids derived from the NH048T tumor sample were cultured without hormone supplementation in the medium and so, *PGR* and *ESR1* expression was lost in culture (Figure S3A-D). Of the three samples – NH85TSc and NH048T have *TP53* driver mutations and *MYC* amplification [46] along with other mutations in cancer driver genes as summarized in Table S2. DS115T displayed a low mutational burden based on both organoid profiling and patient clinical testing (Table S2). However, this patient eventually succumbed to metastatic disease.

**Figure 4:**
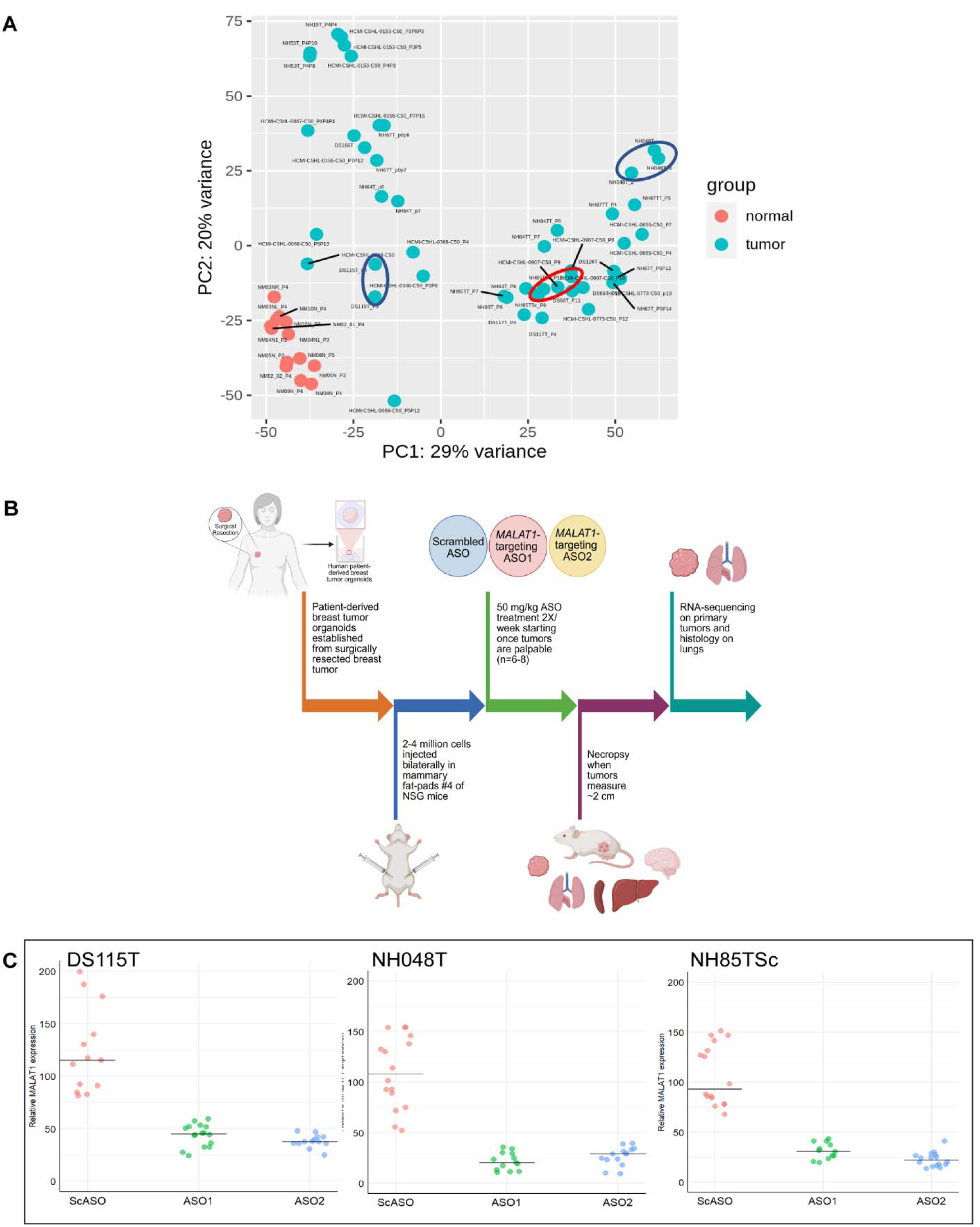
Differential gene expression upon antisense oligonucleotide (ASO)-mediated *MALAT1* knockdown (KD). (A) Principal component analysis (PCA) plot for bulk RNA-seq data of *in vitro* cultured PDO models highlighting the differences between three selected models (circled) used for PDO-X ASO KD experiments. Normal: normal breast organoids derived from tissues post reduction mammoplasty surgeries. Blue circles - Different passages of DS115T and NH048T PDOs each; red circle: replicates of NH85TSc tumor PDOs along with replicates of HCMI-CSHL-0907-C50 i.e. PDOs derived from axillary lymph node recurrence of same patient. (B) Schematic explaining the methodology for development of PDO-X and *in vivo* ASO treatment (Created in https://BioRender.com). (C) Relative *MALAT1* expression level in PDO-X tumors upon necropsy (left) as measured via qRT-PCR, each dot represents one tumor (n=6-8 mice/group)

We performed single molecule RNA Fluorescence in situ hybridization (smRNA-FISH) on the three patient tumor tissues and found *MALAT1* to be upregulated in the tumor cells compared to the adjacent stromal cells (Figure S4). This observation agrees with previously published findings in other patient primary breast tumor tissue samples [14,36]. The three PDO models do not exhibit amplifications in the *MALAT1* chromosomal region (GRCh38/hg38 chr11:65,499,045-65,506,516), as previously reported [46]. Further, to comprehensively characterize the *MALAT1* gene in the three PDO models and to examine if upregulated *MALAT1* in tumor cells can be potentially attributed to mutations or methylation changes, we performed Affinity-based Cas9-Mediated Enrichment (ACME) of specific targets followed by long-read Oxford Nanopore sequencing [63]. Upon examining the mutations and methylation pattern of the *MALAT1* gene body, promoter and putative upstream enhancer region, we found no tumor-specific single nucleotide variants (SNVs) in all three regions in all three PDO models, except one SNV (rs79910129: G→A at GRCh38/hg38 chr11:65,501,008) in the *MALAT1* gene body in NH048T organoids (Table S7). We also sequenced the TP53 gene and found one SNP in exon 7 in the NH048T model (rs28934575: C→T at GRCh38/hg38 chr17:7674230) and a SNP in exon 8 in the NH85TSc model (rs28934576: C→T at GRCh38/hg38 chr17:7673802).

### Systemic ASO delivery led to robust *in vivo* knockdown of *MALAT1* in PDO-X tumors

We chose three distinct patient-derived TNBC organoid models for assessing the impact of *MALAT1* perturbation *in vivo*. For each of the three models, patient-derived breast tumor organoids (PDOs) were injected bilaterally into mammary fat-pad #4 of NOD *scid* gamma (NSG) immunocompromised mice at a density of 2-4 million cells per fat-pad (Figure 4B). The exact number of injected cells depended on the relative *in vitro* growth rate of the organoid model, with 2 million cells/injection for NH85TSc and 4 million cells/injection for DS115T and NH048T PDOs. Once tumors became palpable (∼100 mm^3^), mice were treated subcutaneously with an antisense oligonucleotide (ASO) – either a control non-targeting scrambled ASO (ScASO) or one of two independent *MALAT1*-targeting ASOs (ASO1 or ASO2) at 50 mg/kg, administered twice weekly. Treatment was continued until the ethical endpoint of a maximum palpable tumor diameter of 2 cm, after which necropsies were performed to collect primary tumors (for histology and RNA sequencing), lungs, liver, spleen, kidneys, and brain (Figure 4B).

Bilateral injection of 4 million cells derived from DS115T organoids into mammary fat-pad#4 of NSG mice led to measurable tumors in 2.5 weeks post transplantation. Mice were then randomized into three treatment groups (n=8 per group) and ASO treatment was initiated after tumor establishment and continued until the ethical endpoint. For DS115T PDO-Xs, the ASO treatment duration ranged from 9 to 11.5 weeks long.

Subcutaneous ASO delivery led to a strong knockdown in the primary tumors with an average 59 % *MALAT1* knockdown compared to control group, as measured via qRT-PCR (Figure 4C). Bulk RNA-sequencing was performed on the RNA from the primary tumors. Comparison of *MALAT1* transcript counts per million across treatment groups (Figure S5A) further validated the robust *MALAT1* knockdown observed by qRT-PCR. Differential gene expression analysis revealed 319 human genes altered (p-value<0.01) in DS115T PDO-X tumors upon *MALAT1* knockdown. Of these 319 genes, 162 genes were consistently down-regulated and 157 genes were up-regulated in both ASO groups. *MALAT1* showed a strong >2-fold down-regulation via RNA-seq (Figure S6A) upon treatment with each targeting ASOs. There was no significant reduction in average tumor volume upon *MALAT1* knockdown (Figure S7A).

Next, using the same approach, we generated NH048T PDO-Xs by injecting 4 million cells per fat-pad for transplantation. This model also displayed slow *in vivo* growth and formed measurable tumors at 9.5 weeks post-implantation. ASO treatment was then commenced at 50mg/kg, administered twice a week for 6-14 weeks until the endpoint. Tumor growth rates were highly variable across animals as is evident from the range of treatment duration (Figure S7B). Subcutaneous delivery of ASOs in NH048T PDO-Xs led to the strongest *in vivo MALAT1* knockdown with an average 76% reduction in the primary mammary tumors compared to the control group, as measured via qRT-PCR (Figure 4D). Bulk RNA-sequencing was performed on the RNA from the primary tumors. *MALAT1* knockdown was further validated by RNA-seq (Figure S5B) and tumor volumes were not altered upon knockdown (Figure S7B). Differential gene expression analysis identified 28 human genes significantly (p-value<0.01) altered in NH048T PDO-X tumors upon *MALAT1* knockdown, including 9 downregulated and 19 upregulated genes (Figure S6B). Similar to NH85TSc PDO-Xs, *MALAT1* was about 4-fold down-regulated in NH048T PDO-Xs (Figure S6B).

For generating NH85TSc PDO-Xs, 2 million cells were injected per fat-pad for transplantation as this PDO model grows faster than DS115T and NH048T *in vitro*. However, this model had an intermittent initial take-rate *in vivo* and formed palpable tumors in 5 weeks and the tumors grew quickly once established. ASO treatment (n=6-8 per group) was delivered at 50mg/kg twice a week and continued for 6.5-9 weeks until the ethical endpoint. Strong *MALAT1* knockdown averaged 73% in the primary tumors compared to the control ScASO group, as measured via qRT-PCR (Figure 4E) and was confirmed with ∼4-fold *MALAT1* reduction by bulk RNA-seq (Figure S5C and S6C). No change in tumor volume was observed upon *MALAT1* KD (Figure S7C). 952 human genes were found to be significantly differentially expressed (p-value<0.01) in NH85TSc PDO-X tumors upon *MALAT1* knockdown. Of these 952 genes, 416 genes were consistently down-regulated and 532 genes were up-regulated in both ASO groups. While all three PDO-X models show differential gene expression upon *MALAT1* knockdown, the change in expression for most downstream hits are rather mild as evident from the fold change. The gene expression changes were also found to be patient-specific with no overlap between models, with the exception of *MALAT1*.

### *MALAT1* regulates splicing of a subset of cancer-associated transcription factor targets via regulating intron retention of novel isoforms

To examine if knocking down *MALAT1* leads to changes in alternative splicing, the RNA-seq data was processed through the Envisagenics SpliceDuo pipeline [61]. The splicing changes were categorized into one of four splicing event types (Figure 5A) – alternative acceptor (AA), alternative donor (AD), cassette exon (CA) or intron retention (IR). Cumulatively, 469 differential splicing events (corresponding to 415 genes) were observed (|dPSI|≥0.02) upon knocking down *MALAT1* in DS115T PDO-X tumors by one of two *MALAT1* targeting ASOs vs ScASO control (Figure 5B). NH048T PDO-Xs showed 537 differentially spliced isoforms (corresponding to 516 genes) and NH85TSc PDO-Xs showed 1742 differential splicing events (corresponding to 1413 genes) upon *MALAT1* depletion (Figure 5B). Each of the three independent PDO-X models displayed splicing changes representative of all four types of splicing events (Figure 5B).

**Figure 5:**
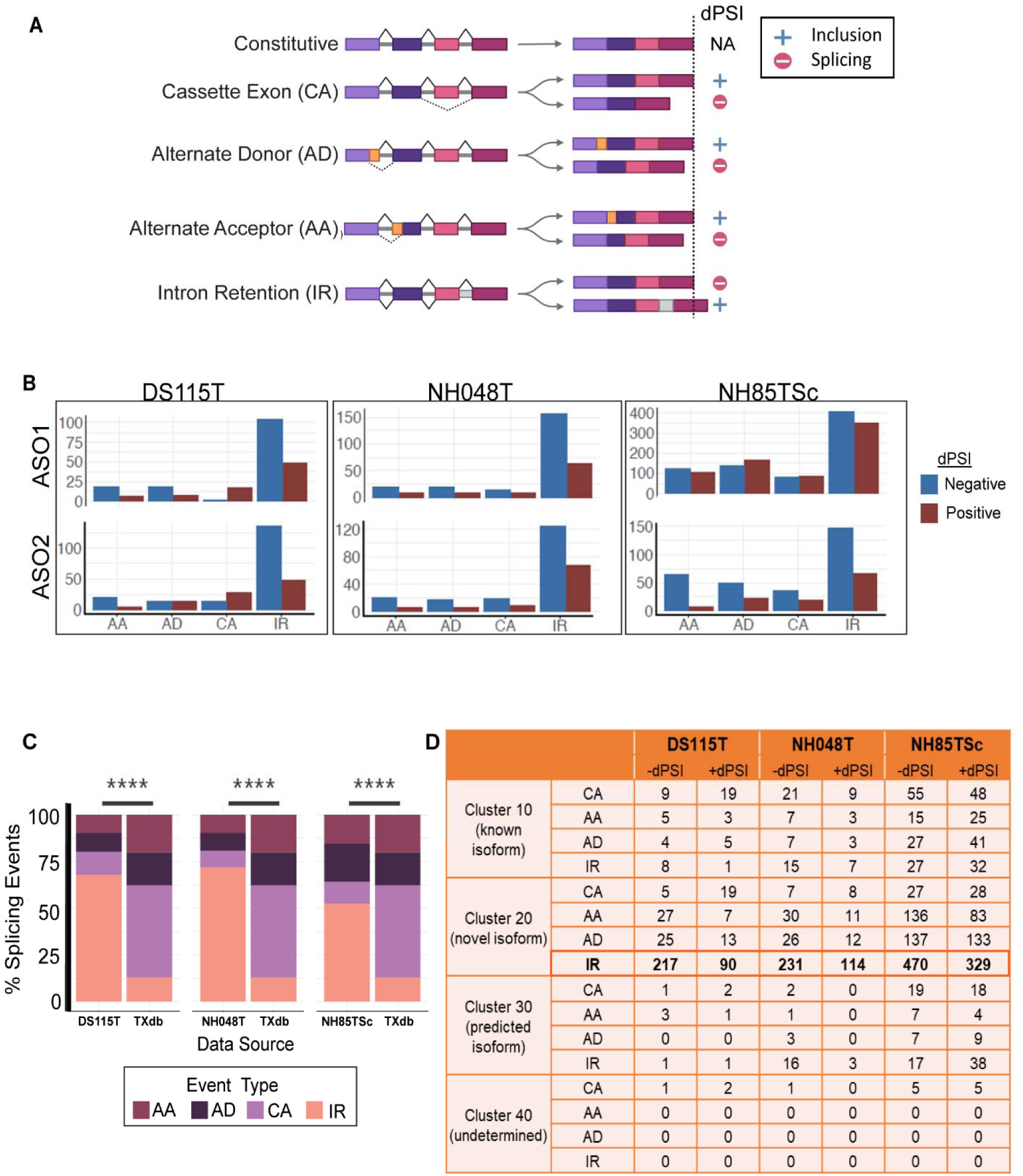
*MALAT1*-dependent changes in alternative splicing in patient-derived breast cancer organoid xenograft (PDO-X) tumors. (A) Schematic for types of alternative splicing event types and the delta percent spliced in (dPSI) direction (Created in BioRender. Aggarwal, D. (2025) https://BioRender.com/evbcrz5). (B) Total number of statistically significant (|dPSI|>0.2, % reproducibility>33%, consistency of dPSI direction>90%) alternative splicing events for each *MALAT1*-targeting ASO (ASO1 and ASO2) compared to scrambled ASO, for each of the 3 PDO-X models – DS115T, NH048T and NH85TSc categorized as per splicing event types. AA: alternate acceptor, AD: alternate donor, CA: cassette exon, IR: intron retention. (C) Percent distribution of splicing event types for each PDO-X model treated with either of *MALAT1*-targeting ASO compared to database (TXdb) of all known splicing events. Statistical analysis using Chi-squared test (****p-value<0.0005). (D) Distribution of splicing events across clusters as defined in the Envisagenics transcript database using the SpliceDuo algorithm. Cluster 10: curated known splicing events, Cluster 20: exon/intron structure is annotated in Ensembl or Ref-seq but isoforms are not reported, Cluster 30: computationally predicted alternative splicing, Cluster 40: predicted gene exon/intron structure. +dPSI indicates increased inclusion and -dPSI indicates increased skipping in *MALAT1*-targeting ASO group compared to control

We discovered that 50-75% of all differential splicing events that occurred upon *MALAT1* knockdown in each of the three PDO-X models involved intron skipping (+dPSI IR) or intron retention (-dPSI IR) irrespective of the PDO-X model (Figures 5C and D). In depth analysis demonstrated that IR events accounted for 318/469 (67.8%) events in DS115T, 386/537 (71.9%) in NH048T, and 913/1,742 (52.4%) in NH85TSc models upon *MALAT1* knockdown (Figure 5D). To verify that this distribution is specific to *MALAT1* perturbation and not a bias resulting from the inherent distribution within the Envisagenics database, the splicing event distribution trend was compared to that of the transcript database (TXdb) of all known splicing events. For each PDO-X experiment, this shift in distribution of splicing changes towards IR events was observed to be statistically significantly different from the inherent distribution of the event types within the database (Figure 5C), indicating a *MALAT1*-specific enrichment of IR events.

Stratification of alternative splicing events by transcript annotation clusters revealed that the majority of the *MALAT1*-dependent splicing changes, irrespective of the PDO-X model, belong to cluster 20, i.e. the exon/intron structure of these genes are annotated in Ensembl and/or Refseq databases, however these isoforms have not previously been reported (Figure 5D). This cluster 20 consisting of unannotated splicing isoforms represented 403/469 (85.9%), 439/537 (81.7%) and 1,343/1,742 (77.1%) of all alternative splicing events in DS115T, NH048T, and NH85TSc, respectively. Within this cluster, intron retention remained the dominant event class, indicating that *MALAT1* depletion preferentially alters intron usage within unannotated transcripts across independent patient-derived tumor models. We observed both +dPSI and -dPSI events within cluster 20 IR group demonstrating that *MALAT1* depletion bidirectionally modulates intron retention. Within cluster 20, -dPSI IR events (217 events in DS115T, 231 events in NH048T and 470 events in NH85TSc) reflect reduced intron inclusion upon *MALAT1* depletion, compared to unannotated isoforms seen in the control group. The +dPSI IR events (90 events in DS115T, 114 events in NH048T and 329 events in NH85TSc) represent increased intron inclusion thereby generating novel unannotated isoforms with retained introns (RI) upon *MALAT1* knockdown (Figure 5D).

Furthermore, we found that the genes corresponding to *MALAT1*-driven splicing changes in the PDO-X tumors associate with a common set of transcription factors across all three independent PDO-X models, as evident from published ChIP-seq data (Figure 6A and B). The numbers in the Venn diagram (Figure 6A) for each PDO-X model represent the TFs observed to be common between ASO1 and ASO2 individually compared to ScASO. Examining the associated transcription factors that were significantly enriched (p-value<0.05) in each of the three PDO-X models, we observed 15 transcription factors to be shared among the three independent models (Figure 6A and B). This subset of specific transcription factors was found to be significantly associated with cancer progression and transcriptional misregulation in cancer (Figure 6C).

**Figure 6:**
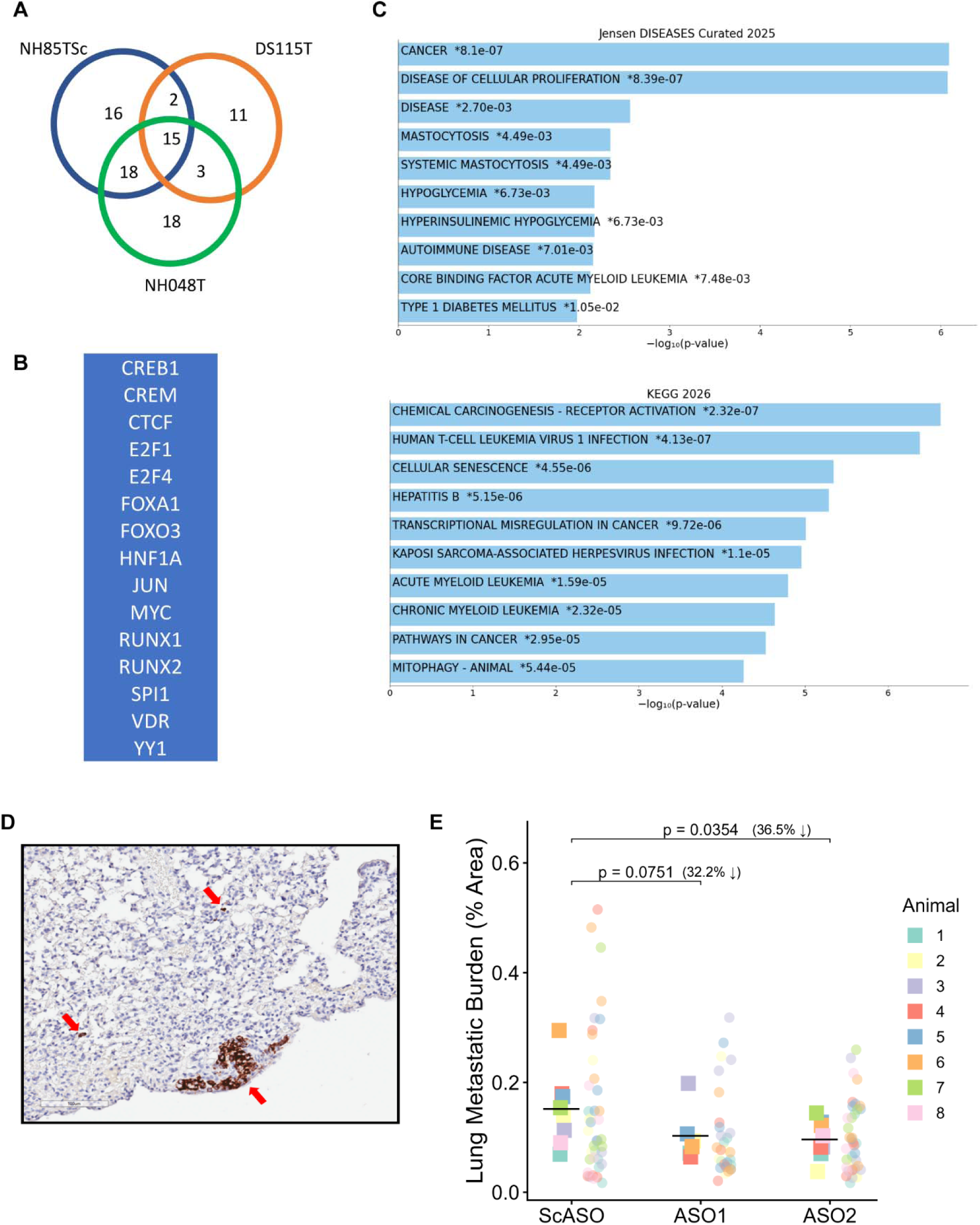
Alternative splicing and phenotypic changes driven by *in vivo MALAT1* knockdown (KD) in PDO-Xs. (A) Venn diagram showing the overlap between the number of transcription factors significantly associated with alternatively spliced transcripts upon *MALAT1* KD as per the ChEA 2022 ChIP-seq database on Enrichr (p-value<0.05) for each of the three PDO-X models. (B) List of 15 significantly associated transcription factors common between the three PDO-X models (p-value <0.05). (C) The ontologies and pathways enriched for the list of 15 shared transcription factors as per Jensen Diseases Curated 2025 database and KEGG 2026 Human database via Enrichr. (D) Representative area of a 5µm lung section from a NH85TSc PDO-X animal treated with ScASO, immuno-stained with anti-human mitochondria antibody (brown), nuclei (blue). Red arrows indicate areas of positive staining. (E) Lung metastatic burden as average % lung area stained, in NH85TSc PDO-Xs as measured via QuPath for five serial sections 100 µm apart from each other per lung (n=6-8/group) represented as dots and average of all sections per animal represented as squares, horizontal black line represents mean, p-value as measured via Welch’s one-tailed t-test assuming unequal variances. ScASO – scrambled ASO; ASO1 and ASO2 – *MALAT1* targeting ASOs.

### *MALAT1* depletion reduces lung metastatic burden in TNBC PDO-X models

Given the previously reported 70% reduction in lung metastasis upon knocking down *MALAT1* in the MMTV-PyMT mouse model of breast cancer [14], we wanted to assess any potential changes in metastasis in the PDO-X models upon *MALAT1* knockdown. However, PDO-X tumors do not readily metastasize, whereas some PDO-X models form single cell metastases or micro-metastases in the lungs (Figure 6D, red arrows). Since there are no large metastatic nodules developed unlike mouse models of breast cancer, it is particularly challenging to quantitate metastases in these advanced PDO-X models. Using a previously published IHC protocol employing an antibody targeting human mitochondria [46], we were able to identify these single cell and micro-metastases which were then quantitated using QuPath software. Overall, less than 1% of the total lung area in a given section contained metastases. The quantification showed high variability between sections taken from different depths of the lung tissue blocks, each of which had lungs embedded as separated lobes. An average of the quantification across 5 sections (each 5 µm thick) that were each 100 µm apart within the block for each animal per treatment group (n=6-8 mice/group) showed an average 32.2% decrease in metastasis upon knockdown with ASO1 and an average 36.5% decrease with ASO2 (Figure 6E).

### Knocking down *MALAT1* in PDO-X tumors impacts the abundance of macrophages in the tumor microenvironment

Next, since the PDO-X tumors were infiltrated by mouse stromal cells, all RNA-seq data was aligned to a combined reference genome of human and mouse origin. The human and mouse gene counts were split before DESeq2 analysis to avoid bias. On aligning the RNA-seq data to the mouse reference genome, we observed 797 differentially expressed mouse genes common between each of the *MALAT1*-targeting ASOs compared individually to ScASO in NH85TSc PDO-Xs (Figure 7A and B). For NH048T PDO-Xs, 563 differentially expressed mouse genes common between the two ASOs (Figures 7A and B). Further, we observed 587 differentially expressed mouse genes common between the two ASOs in DS115T PDO-Xs (Figures 7A and B). It must be noted that the mouse homolog of *Malat1* was not differentially expressed in the RNA from stromal cells (Figure S8A-C) in all three PDO-X models, thus confirming that the ASOs bind specifically to human *MALAT1*.

**Figure 7:**
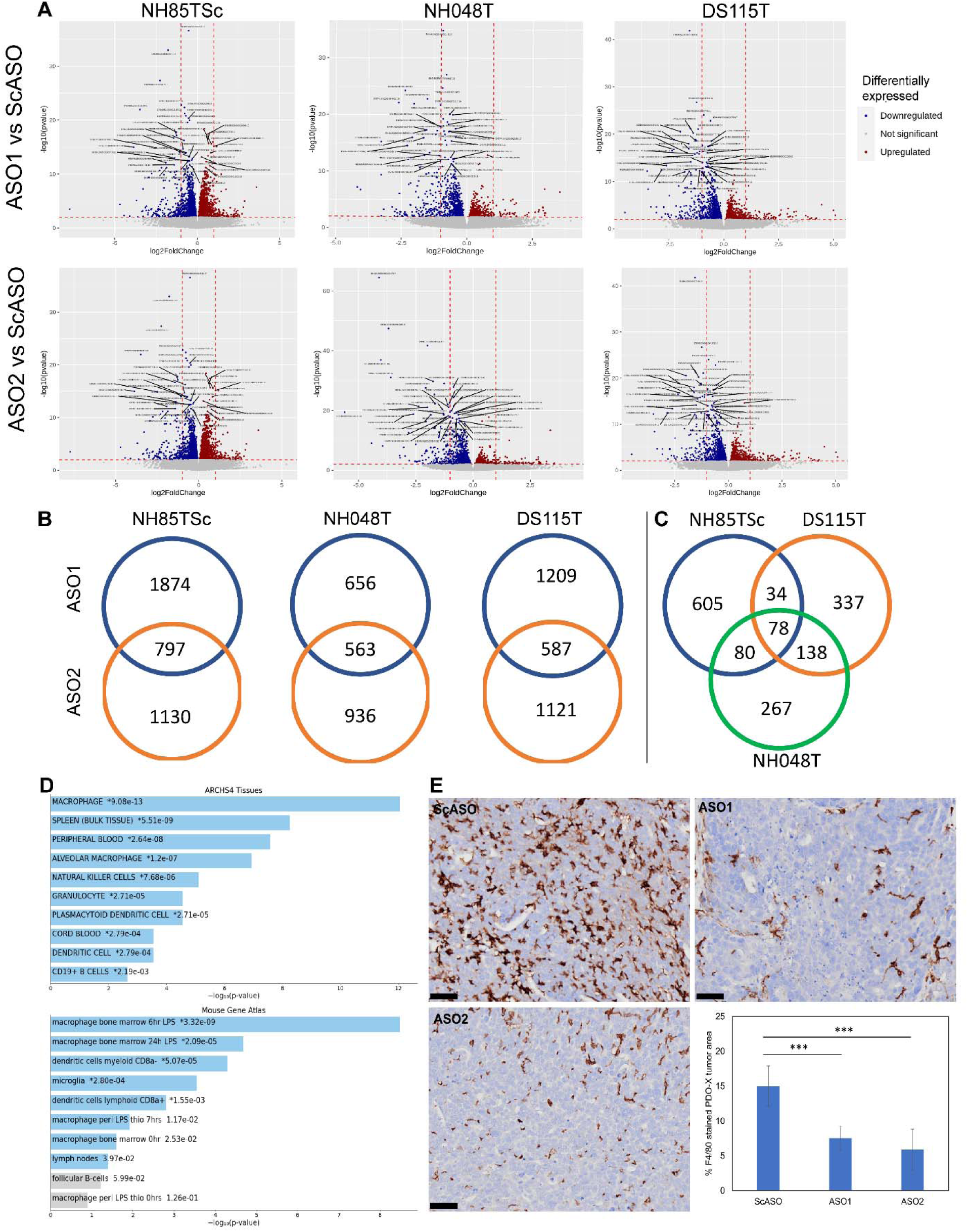
*MALAT1*-dependent changes in the tumor microenvironment. **(A)** Volcano plots displaying differentially expressed genes in the mouse stromal cells for each of the three PDO-X models. Red dots: upregulated mouse genes upon human *MALAT1* knockdown (KD) in tumor cells, blue dots: downregulated mouse genes. (B) Venn diagrams showing overlap of differentially expressed (up and downregulated) mouse genes for each *MALAT1*-targeting ASO (ASO1 and ASO2) compared to scrambled ASO. (C) Venn diagram showing overlap of differentially expressed mouse genes between three PDO-X models. (D) Cell types associated with common downregulated mouse genes upon *MALAT1* KD as per ARCHS4 and Mouse Gene Atlas databases. (E) Immunohistochemistry representative images showing labeling of NH85TSc PDO-X tumors with anti-mouse F4/80 antibody and corresponding quantitation of F4/80 positive area examining macrophage infiltration within tumor sections, via QuPath using the mean of >3 representative areas (∼1mm^2^) from each tumor section (n=6 tumors/group). ScASO – scrambled ASO; ASO1 and ASO2 – *MALAT1* targeting ASOs. Scale bar = 50 µm. Error bars represent standard deviation. *** P<0.001 using the two-tailed unpaired Welch’s t-test.

Upon assessing the overlap, we found 78 genes to be common among the differentially expressed mouse genes from all three organoid lines (Figure 7C). Of the 78 genes, 74 mouse genes were consistently downregulated in all the individual PDO-X experiments whereas only 4 mouse genes were found to be upregulated upon *MALAT1* knockdown. Pathway analysis of the common differentially downregulated mouse genes among the three different PDO-X models upon *MALAT1* knockdown compared to ScASO-treated animals was performed using the Enrichr web tool. The Reactome 2024 pathways database revealed these differentially downregulated mouse genes to be associated with various immune pathways such as immune system, innate immune system, neutrophil degranulation, cytokine signaling in immune system etc. (Figure S9). Interestingly, evaluating the cell-types represented by this subset of common downregulated genes revealed a strong association with macrophages and monocytes. This was consistently observed across multiple cell-type databases available via Enrichr, two of which (Mouse Gene Atlas database and ARCHS4 databases) are shown in Figure 7D.

We further examined the expression of the macrophage-specific marker F4/80 via immunohistochemistry. Sections from six tumors (n=3 mice/treatment group) with NH85TSc PDO-Xs were labeled using the monoclonal antibody targeting F4/80 (Figure 7E). The data was quantified using QuPath software’s pixel classifier feature identifying DAB staining on a hematoxylin background stain. We observed a 7% decrease in the abundance of tumor-infiltrating macrophages (F4/80 positive within dense tumor regions) upon ASO1 knockdown and a 9% decrease upon ASO2 knockdown, compared to ScASO (Figure 7E).

## Discussion

In this study, we systematically evaluated *MALAT1* knockdown efficiency across a large panel of 28 human patient-derived breast tumor organoid (PDO) models representing various subtypes of breast cancer using two independent antisense oligonucleotides (ASOs). Consistent with prior work targeting *KRAS* using ASOs in diverse cancer cell line models [70], we did not observe a correlation between endogenous *MALAT1* abundance or RNaseH1 expression and ASO potency in these patient-derived cancer models. Further, we also confirmed that neither ASO penetration within the 3D Matrigel/organoid domes nor cellular uptake are limiting factors influencing knockdown efficiency. Although, subcellular distribution and protein interactions are known to impact ASO efficiency [70,71], we did not assess these factors here. Notably, fast-growing PDO models often derived from high-grade, poorly differentiated tumors consistently displayed the most robust ASO-mediated *MALAT1* knockdown *in vitro*. Given that the majority of these aggressive PDO models were derived from TNBC tumors, these findings suggest that a *MALAT1*-targeting ASO therapy may be particularly effective in treating aggressive TNBC tumors in the clinic.

Building on these *in vitro* findings, we optimized an organoid engraftment strategy and successfully established and characterized ten patient-derived organoid xenograft (PDO-Xs) models. We observed that PDO-X tumors grow at widely variable rates and did not necessarily mirror the growth kinetics observed *in vitro*, highlighting the influence of the *in vivo* microenvironment. PDO-Xs are excellent models as they are clinically relevant and preserve key patient-specific tumor features while mitigating practical limitations of patient-derived xenografts (PDXs) such as scalability, dependency on amount of starting material, turnaround time, rate of distant metastases etc. [72,73]. Using three independent breast tumor PDO-X models – NH85TSc, NH048T and DS115T, systemic subcutaneous delivery of *MALAT1-*targeting ASOs achieved robust knockdown efficiencies ranging from 59-76% *in vivo*. Transcriptomic analyses following *MALAT1* depletion revealed relatively modest changes in gene expression within primary tumors, suggesting that *MALAT1* may act as a regulator fine-tuning transcriptional programs rather than a binary switch. Alternatively, *MALAT1*-dependent transcriptional effects may be restricted to specific tumor sub-populations, with bulk RNA-seq averaging these effects.

Interestingly, the transcriptional changes we observed were rather modest and highly patient-specific, consistent with previous reports demonstrating context-dependent *MALAT1*-regulated transcriptional changes across various 2D and *in vivo* models [reviewed in 22,23]. TNBC is widely recognized as a highly heterogeneous disease, with inter-patient variability extensively characterized at multiple levels – histopathological [31], genomic [31,35] and transcriptomic [32–34,39]. Our data similarly revealed pronounced transcriptomic diversity across TNBC PDO models (Figure 4A). Beyond inter-patient heterogeneity, TNBC tumors also display substantial intra-tumor heterogeneity (ITH), as previously demonstrated using next-generation sequencing and spatial transcriptomics approaches [40–42,42,43]. Examining whether the patient-specific changes in the transcriptome observed here reflects patient variability, intra-tumor heterogeneity, or both will require future analyses beyond the scope of this study incorporating single-cell and spatial transcriptomics approaches.

*MALAT1* is a well-established nuclear speckle-localized lncRNA shown to interact with pre-mRNA splicing factors, nascent transcripts and active gene loci, thereby influencing spliceosome assembly and pre-mRNA processing in 2D cell line and mouse models [14,16,18,19,74–78]. Consistent with a proposed model of *MALAT1* acting as a molecular scaffold orchestrating splicing regulation, we observed widespread changes in alternative splicing upon *MALAT1* depletion in human PDO-X tumors. These changes spanned all four major classes of alternative splicing events, and this observation was reproducible across independent patient-derived tumor xenograft models, supporting a conserved regulatory role for *MALAT1* in determining alternative splicing outcomes *in vivo*.

Notably, upon evaluating the distribution of the four splicing event types among *MALAT1*-driven changes, we observed that intron retention (IR) events represent the major class of splicing alterations in all three PDO-X models. This enrichment was found to be specific to *MALAT1* and not reflective of the baseline distribution within the database of annotated splicing events. While intron retention and splicing were both enabled by *MALAT1* for specific transcripts, we observed a net enrichment of events showing reduced intron retention (-dPSI) following *MALAT1* knockdown. This directional shift is consistent with prior work in mouse embryonic stem cells demonstrating that nuclear *MALAT1* depletion leads to increased splicing of specific introns compared to control, suggesting that *MALAT1* can facilitate intron retention at selected loci [79]. Together, these observations indicate that *MALAT1* modulates intron definition in a locus- and tumor-context-dependent manner, rather than acting as a global regulator of splicing efficiency.

Strikingly, *MALAT1*-dependent IR splicing changes were highly enriched within a cluster of previously unannotated transcript isoforms, indicating that *MALAT1* influences isoform composition primarily through alternative usage of known exon-intron structures. These unreported isoforms, particularly ones with retained introns (RI), generated upon *MALAT1* depletion (i.e. +dPSI), raise potential immunologic implications. Prior work has shown that RI-derived peptides can be presented on MHC-I molecules [80], associate with improved patient survival [81], and promote T-cell activation and tumor cytotoxicity [82,83]. Pharmacologic modulation of splicing has been shown to generate immunogenic neoantigens (neoAgs) that improved endogenous T-cell response to the tumor *in vivo* as well as augmented immunotherapy response [84]. Notably, recent *in vivo* studies have demonstrated enhanced cytotoxic T-cell activity upon *Malat1* depletion in multiple immunocompetent TNBC mouse models [24,25] and reduced cytotoxic lymphocyte infiltration in *Malat1*-overexpressing lung tumors [85]. Although the NSG PDO-X models used in the current study lack adaptive immunity and therefore do not permit direct assessment of immune-mediated inhibition of tumor growth, our findings raise the possibility that intron-retaining isoforms generated upon *MALAT1* depletion represent a potential source of tumor-specific neoAgs that may underlie the reduced immune evasion of tumors, observed in immunocompetent models. These findings warrant future studies integrating splicing-derived ORF prediction with MHC-I immuno-peptidomics and functional assessment in humanized immunocompetent mice to determine whether *MALAT1* knockdown-derived isoforms with retained introns contribute to enhanced immune-mediated tumor cell clearance.

Importantly, genes undergoing *MALAT1*-dependent splicing changes were significantly enriched for targets of 15 shared, specific cancer-associated transcription factors (TFs) across independent PDO-X models. This reproducible convergence of cancer-associated transcription factor networks among *MALAT1*-driven splicing changes across ASOs and distinct PDO-X tumors indicates that *MALAT1*-mediated splicing is functionally coupled to tumor-driving transcriptional circuitry. The 15 shared TFs include canonical proliferation drivers (MYC, E2F1, E2F4)), lineage determinants (FOXA1, RUNX1, RUNX2), chromatin architecture regulators (CTCF, and YY1), and the myeloid lineage regulator SPI1. Notably, some of the shared TFs, particularly MYC, RUNX2, JUN and YY1 have established roles in regulating epithelial-mesenchymal transition, suggesting that *MALAT1*-sensitive splicing events may govern tumor cell plasticity. The consistent overlap of these distinct yet functionally interconnected oncogenic transcriptional modules suggests that *MALAT1*-sensitive splicing events are not distributed randomly across the transcriptome, rather they are embedded within core oncogenic regulatory programs that define cell state, lineage identity and immune interaction.

Beyond tumor-intrinsic effects, we further showed that selective depletion of *MALAT1* in tumor cells without altering *Malat1* expression in the stromal compartment elicited transcriptional changes within the mouse stromal cells. Specifically, 74 genes, associated with macrophage abundance were consistently downregulated upon *MALAT1* knockdown across all three PDO-X models. This aligns with previously published work showing *Malat1*-dependent macrophage infiltration of tumors in mouse models of breast and lung cancer [25,85]. Therefore, the PDO-X model system offers a remarkable advantage in their utility for dissecting certain aspects of tumor-stroma crosstalk *in vivo*.

*MALAT1* has been consistently implicated in promoting metastasis across multiple cancer types, including breast and lung cancer [10,14,24,85,86] and prior work from our laboratory demonstrated ∼70% reduction in lung metastatic burden in the MMTV-PyMT mouse model upon *Malat1* depletion [14]. In the present study, we observed a significant reduction in lung metastases in the NH85TSc PDO-X model following *MALAT1* knockdown. This represents the first quantitative assessment of distant metastasis using human breast tumor PDO-derived xenografts. It is important to note that these human PDOs do not metastasize as readily as cell line-based or genetically engineered mouse models, likely reflecting patient biology, as most breast cancers are non-metastatic at diagnosis [87]. When metastases do arise in the examined PDO-Xs, they are typically limited to single-cell or micro-metastases covering less than 1% of the total lung area in a section, making detection and interpretation particularly challenging. Despite these challenges, the observed 32-36% reduction in metastatic burden is notable and warrants follow-up studies using extended observation post-surgical resection of tumors, and/or tail-vein transplantation of luciferase-labeled PDO models to further validate and extend these findings. Further detailed assessment of the effect of *MALAT1* perturbation on primary tumor growth and metastasis in humanized immunocompetent mice could reaffirm the role of *MALAT1* in crosstalk with the T-cell compartment in human models, as previously observed in syngeneic mouse models of breast cancer [24,25].

In summary, the present study is the first to leverage human patient-derived breast tumor organoid xenografts for a comprehensive, *in vivo* functional interrogation of a long non-coding RNA target. We hereby demonstrate the potential to successfully adopt PDO-X model systems for biological studies beyond drug screening, and we provide a well-characterized database for PDO models suitable for ASO-mediated gene perturbation studies. Our findings conclusively establish *MALAT1* as a key regulator of intron splicing, and its perturbation led to alternative splicing of genes controlled by cancer-associated transcription factors resulting in the generation of novel splicing events. In this context, our splicing data demonstrating novel splicing events suggests an additional mechanistic axis that could explain the role of *MALAT1* in immune evasion of breast cancer cells. Taken together, our data using human patient-derived systems demonstrates that *MALAT1* influences breast cancer progression through coordinated regulation of splicing of cancer-associated transcripts, modulating macrophage abundance in the tumor microenvironment and influencing the metastatic behavior of tumor cells. These findings strengthen the rationale for *MALAT1*-targeting ASOs as a promising therapeutic strategy for breast cancer, particularly the aggressive TNBC subtype, and highlight the value of human PDO-X models for functional and translational studies of RNA-targeted cancer therapies.

### Authors’ Contributions

Conception and design: DA and DLS. Animal work and organoid culture: DA and SR. Data collection: DA, SR, PN, SS, SVI, SB, QG and SP. Data analysis and interpretation: DA, KA, TF, MK, DLS, and JEW. Data analysis support: RU, JAR, GA. Provision of surgical samples and materials: KK, and AR. Oversight of DNA Sequencing: WRM. Oversight of RNA splicing analysis: MA. Funding acquisition: DLS. Writing – DA and DLS. Review and editing – All authors.

## Acknowledgements

We thank members of the Spector lab for critical discussions and advice throughout the course of this study. The authors are deeply grateful to patients and their families for consenting to donate excess tissue for research. We thank the Northwell Health Biorepository and Pathology teams for their efforts and support. The authors thank Dr. Zhen Zhao for guidance on immunohistochemistry analysis, Dr. Tse-Leun Wee for her assistance with confocal microscope use, Dr. Sara Goodwin for technical guidance on long-read sequencing, Jill Habel and Maria Mosquera for assistance with mouse experiments. The authors are grateful to Dr. Adrian Krainer, Dr. Mikala Egeblad, Dr. Zhen Zhao and Dr. Jingfang Ju for their guidance throughout the course of this study.

## Ethics declarations

Initial tumor samples from breast cancer patients were obtained from Northwell Health in accordance with Institutional Review Board protocol IRB-03-012 and IRB 20-0150. The collection of genomic and phenotypic data was consistent with 45 CFR Part 46 (Protection of Human Subjects) and the NIH Genomic Data Sharing (GDS) Policy. Informed consent ensured that the de-identified materials collected, the models created, and data generated from them can be shared without exceptions with researchers in the scientific community. All animal procedures and studies were carried out in accordance with the CSHL Animal Care and Use Committee (IACUC Protocol No. 2021-1197).

## Funding

We acknowledge the CSHL Cancer Center Shared Resources (Animal, Bioinformatics, Next Generation Sequencing, Histology, Microscopy) for services and technical expertise (NCI 2P3OCA45508). The sequencing analysis was performed using equipment purchased through NIH grant S10OD028632. The acquisition of the Zeiss LSM980 confocal microscope with Airyscan2 was funded by NIH 1S10OD034372. This research was supported by NCI 5P01CA013106-Project 3 (D.L.S.) and CSHL/Northwell Health. The content is solely the responsibility of the authors and does not necessarily represent the official views of the National Institutes of Health.

## Abbreviations

AA: alternative acceptor
AD: alternative donor
ACME: Affinity-based Cas9-Mediated Enrichment
AS: Alternative splicing
ASO: antisense oligonucleotide
BC: breast cancer
BSA: bovine serum albumin
CA: cassette exon
cEt: 2’-constrained ethyl
CNV: copy number variation
DEG: differentially expressed gene
dPSI: delta percent spliced in
ER: estrogen receptor
FDA: Food & Drug Administration
FFPE: formalin-fixed paraffin-embedded
H&E: hematoxylin and eosin
HMW: high molecular weight
IDC: invasive ductal carcinoma
IHC: immunohistochemistry
ILC: invasive lobular carcinoma
IR: Intron retention
ITH: intratumor heterogeneity
KD: knockdown
lncRNA: long non-coding RNA
*MALAT1*: Metastasis Associated Lung Adenocarcinoma Transcript 1
MHC-I: Major Histocompatibility Complex class I NeoAgs – Neoantigens
ONT: Oxford Nanopore Technologies
PCA: principal component analysis
PDO: Patient-Derived Organoid
PDO-X: Patient-Derived Organoid Xenograft
PDX: Patient-Derived Xenograft
PR: progesterone receptor
PS: phosphorothioate
qRT-PCR: quantitative real-time polymerase chain reaction
RI: retained introns
RNA: ribonucleic acid
RNA-seq: RNA sequencing
RT: reverse transcriptase
ScASO: Scramble ASO
smRNA-FISH: single-molecule RNA fluorescence in situ hybridization
SNVs: single nucleotide variants
SVs: structural variants
TEM: Transmission Electron Microscopy
TFs: transcription factors
TNBC: triple negative breast cancer
TPM: transcripts per million

## Notes

### Competing Interest Statement

The authors have declared no competing interest.

### Summary of Updates

Various changes have been made throughout the manuscript.

