## Supplementary Figures for "Patient-derived organoid xenografts reveal the multifaceted role of the lncRNA *MALAT1* in breast cancer progression"

**Figure S1**

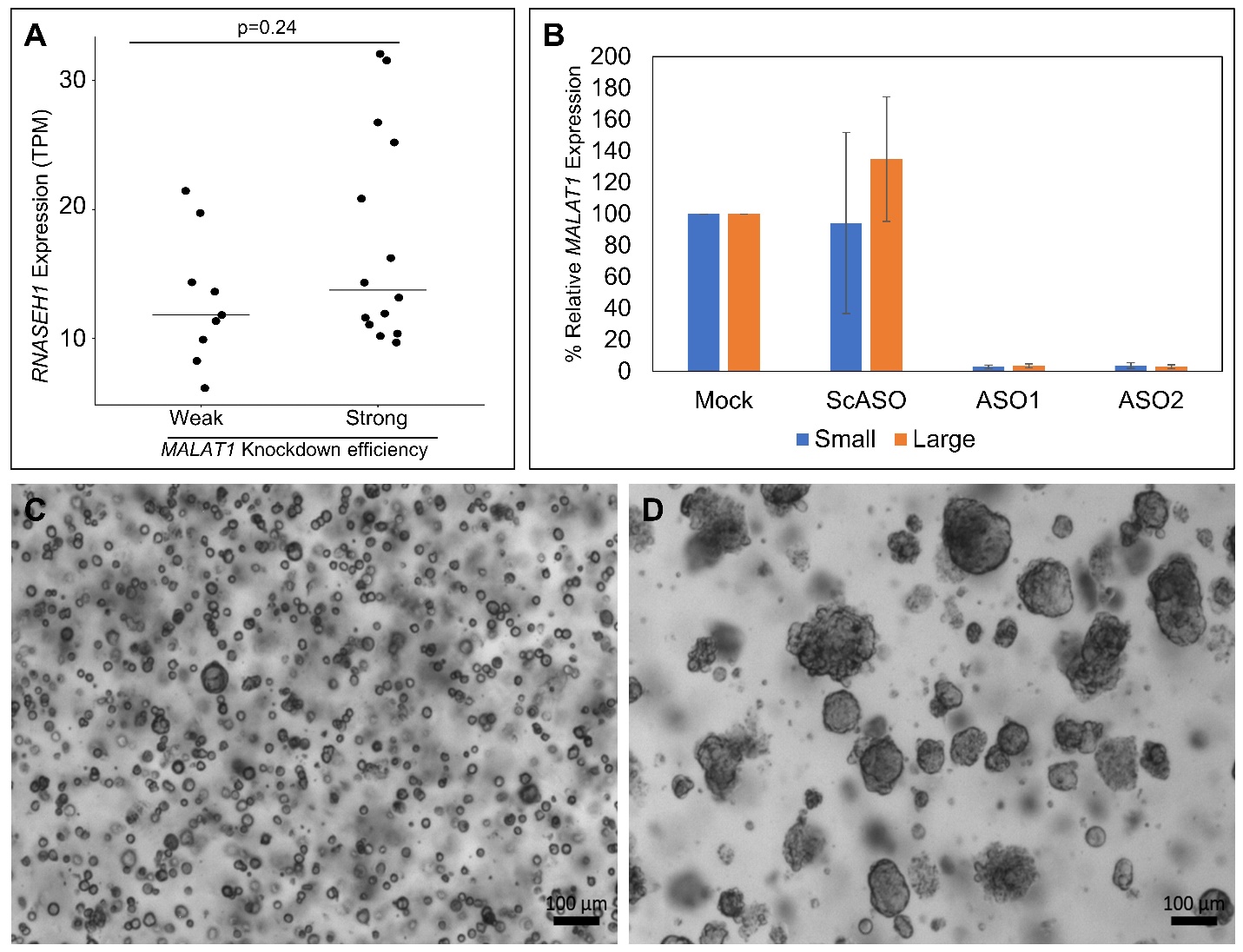

**Supplementary Figure 1: Assessment of *RNASEH1* transcript level and organoid plating size on ASO-mediated *MALAT1* knockdown efficiency.**

(A) *RNASEH1* transcripts per million (TPM) counts from bulk RNA-seq data comparing patient-derived organoid (PDO) models that gave weak vs strong ASO-mediated *MALAT1* knockdown efficiency. P-value calculated using Wilcox test. (B) *MALAT1* knockdown efficiency of two independent ASOs post 6 days of ASO treatment in the PDO models HCMI-CSHL-0366-C50 when plated as small/large organoids on day0 (3µM ASO, n=3). (C-D) Brightfield representative image displaying HCMI-CSHL-0366-C50 PDOs plated as small (C) or large (D) organoids on day 0 prior to ASO treatment.

**Figure S2**
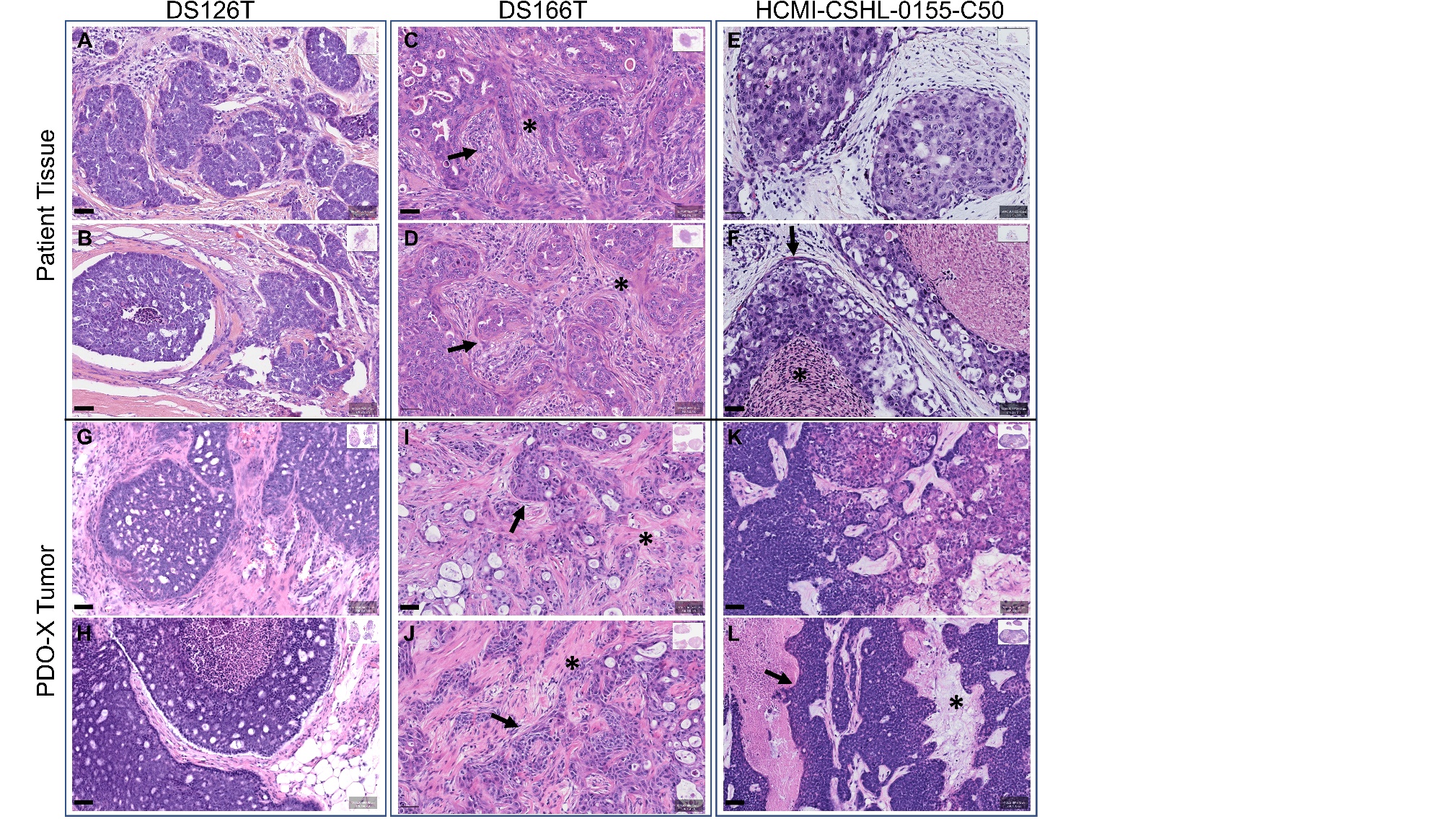

**Supplementary Figure 2: Morphologic comparison of original breast cancer patient tumors to matched patient-derived organoid xenograft (PDO-X) tumors.**

Hematoxylin and eosin (H&E) staining of original patient tumor (A-F) slides and corresponding PDO-X tumor (G-L) in NSG mice for DS126T, DS166T and HCMI-CSHL-0155-C50 PDO models. Scale bar = 50 µm.

**Figure S3**

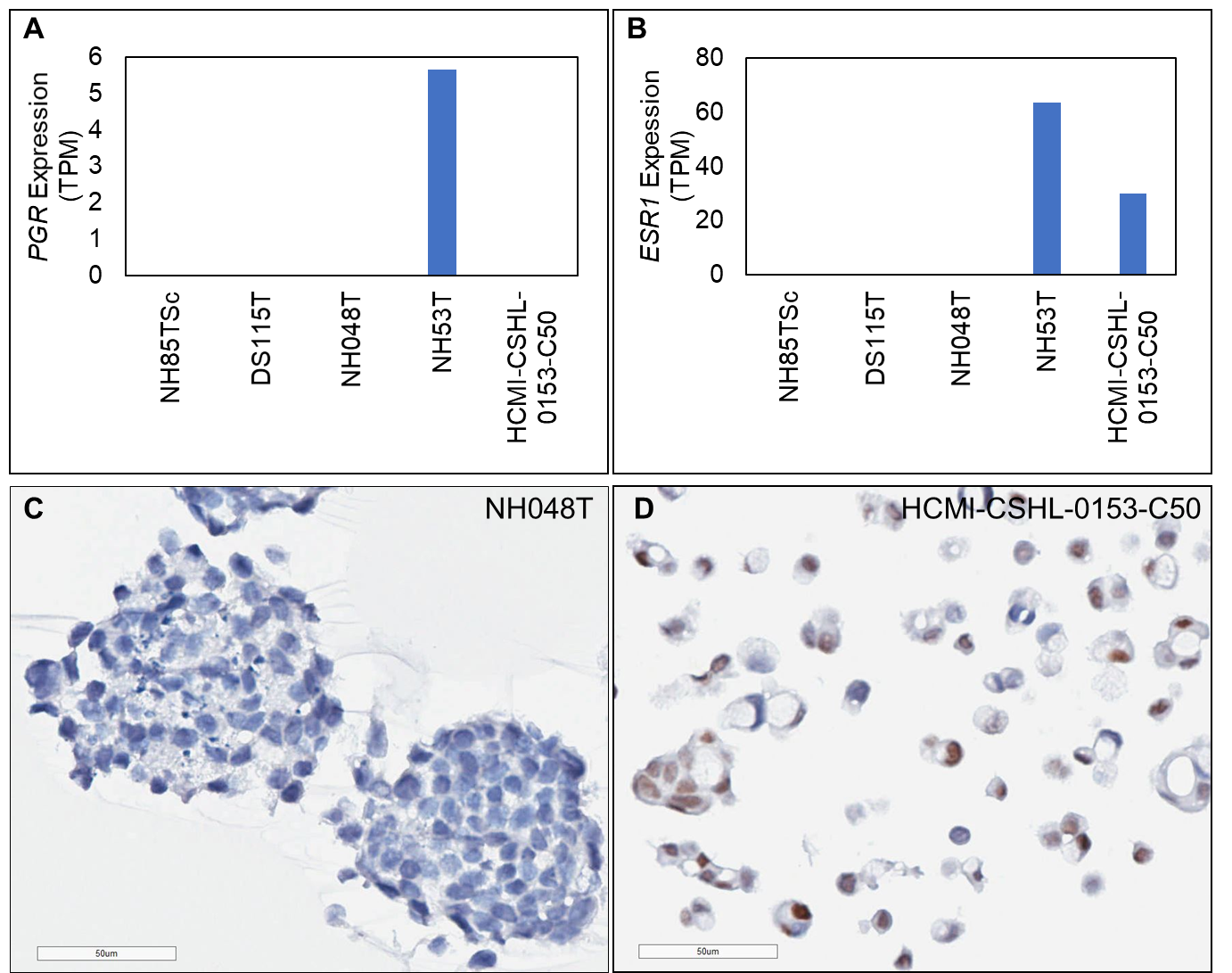

**Supplementary Figure 3: Assessing expression of ER-alpha at the transcript and protein level.**

(A) *PGR* and (B) *ESR1* transcripts per million as per bulk RNA-seq data for the three TNBC PDO models used for PDO-X study. NH53T is an ER+ luminal tumor-derived model and HCMI-CSHL-0153-C50 is an ER+ invasive lobular carcinoma-derived model. (C-D) Immunohistochemistry staining with an antibody targeting ER-alpha (brown, nuclear) with a hematoxylin (blue) counterstain for nuclei in NH048T (C) and HCMI-CSHL-0153-C50 PDOs (D). Scale bar = 50 µm.

**Figure S4**
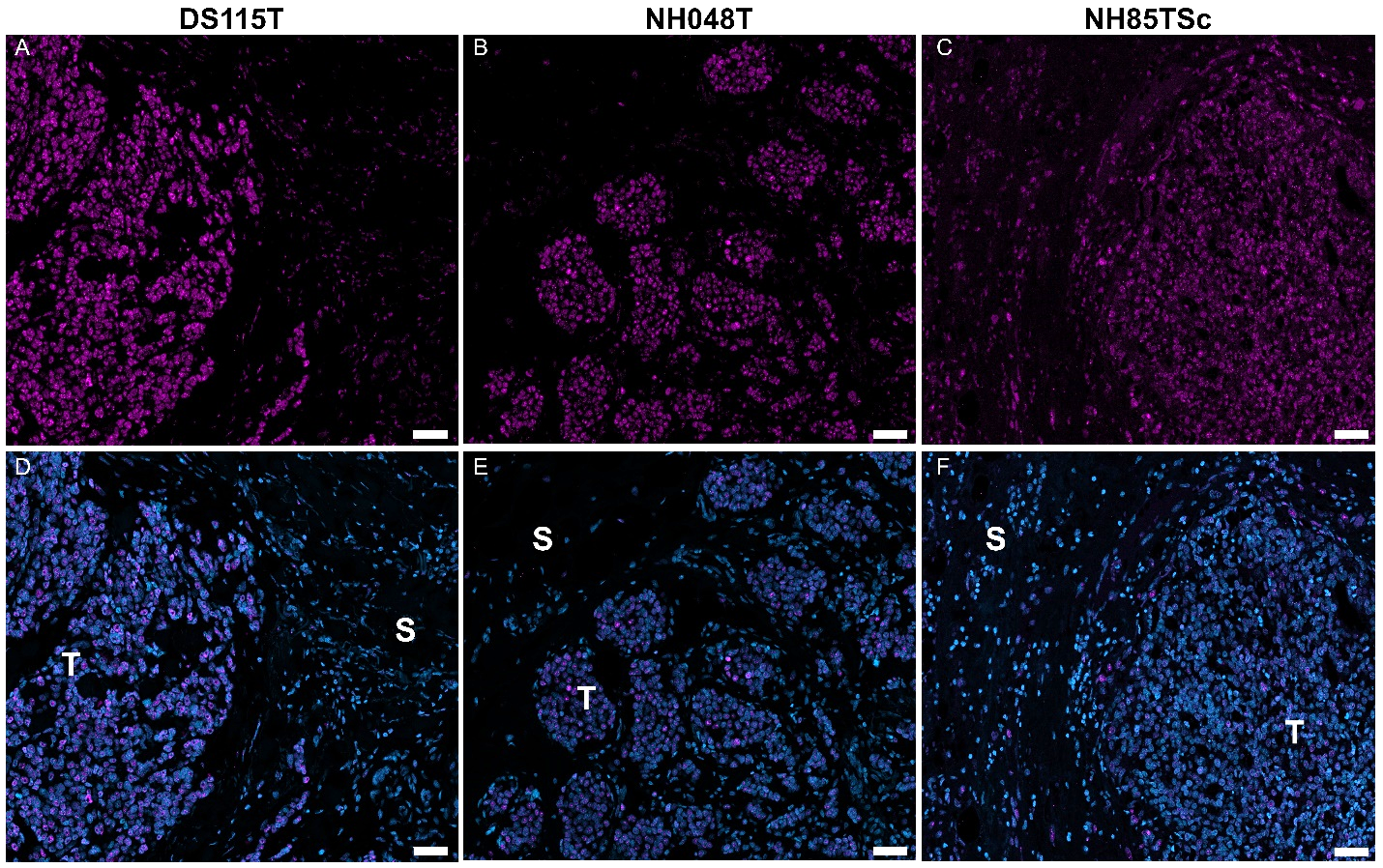

**Supplementary Figure 4: *MALAT1* RNA level in patient tumor sections.**

(A-C) *MALAT1* (pink) expression as visualized by single-molecule RNA fluorescence in situ hybridization on patient breast tumor tissue sections post a surgical resection from the three patients – DS115T, NH048T and NH85TSc. (D-F) Merged channel imaged with *MALAT1* (pink) and DAPI-stained nuclei (blue). S- stroma, T-tumor lesions, scale bar=50µm.

**Figure S5**
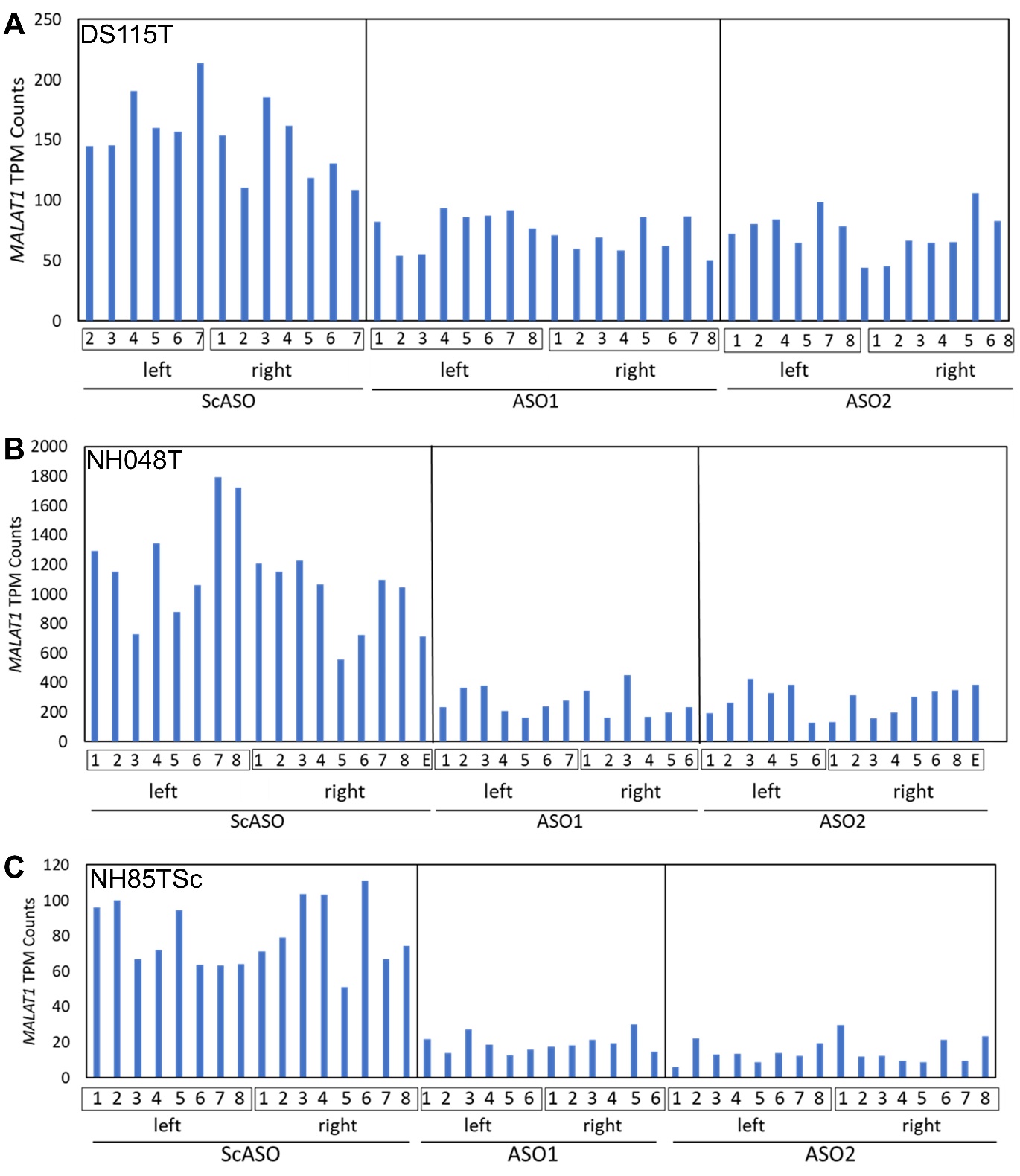

**Supplementary Figure 5: *MALAT1* transcript counts as measured via bulk RNA-seq in PDO-X-derived tumors.**

*MALAT1* transcripts per million (TPM) counts for individual PDO-X-derived tumors for DS115T, NH048T and NH85TSc PDO-Xs treated with either ScASO/ ASO1/ ASO2. Each number corresponds to animal # and left/right represent the fat-pad location. E - additional separate tumor at one of the locations. ScASO – Scramble ASO control, ASO1 and ASO2 – *MALAT1*-targeting ASOs.

**Figure S6**

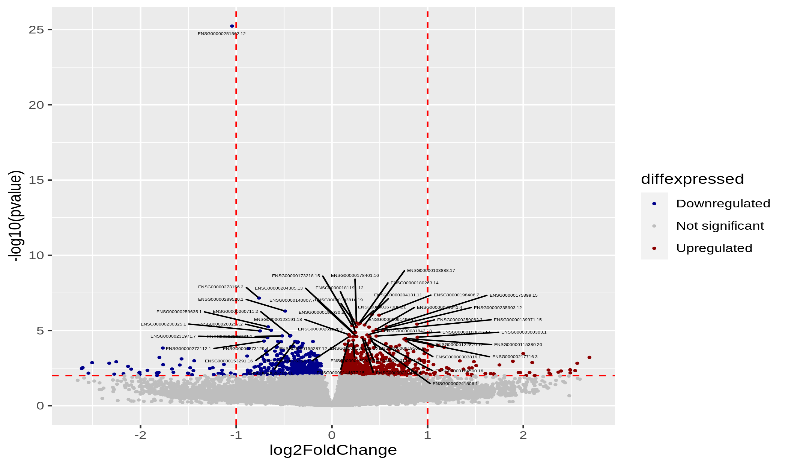

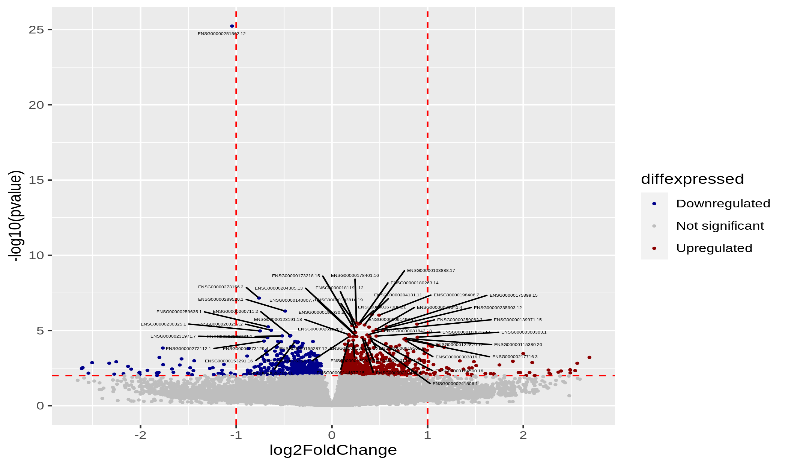

ASO1 vs ScASO

ASO2 vs ScASO

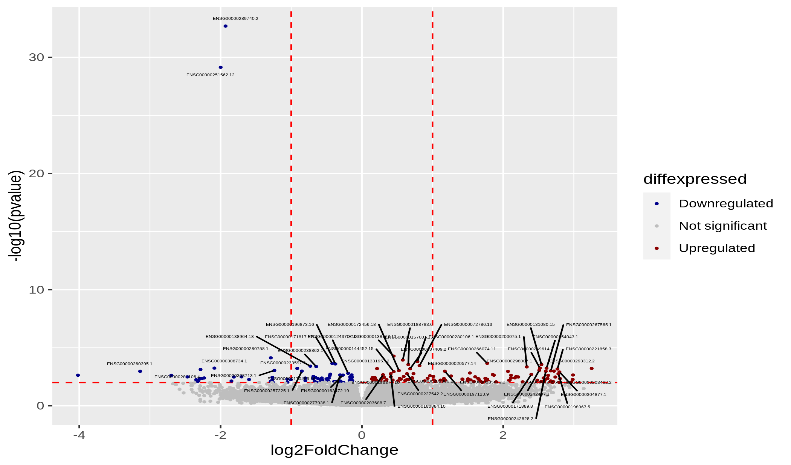

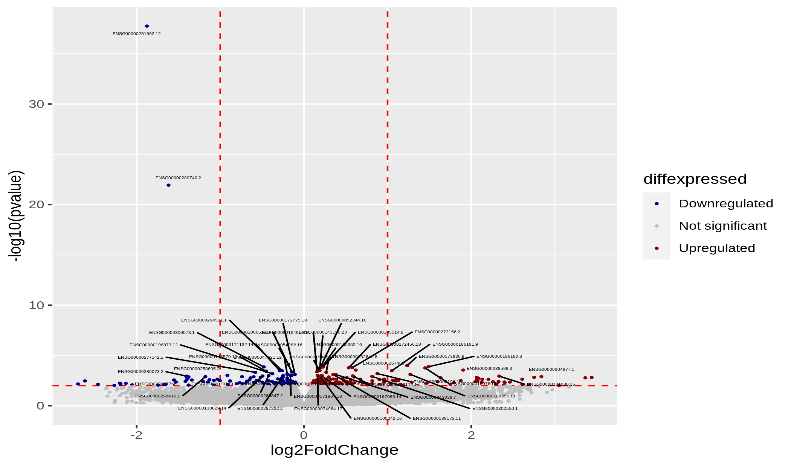

ASO1 vs ScASO

ASO2 vs ScASO

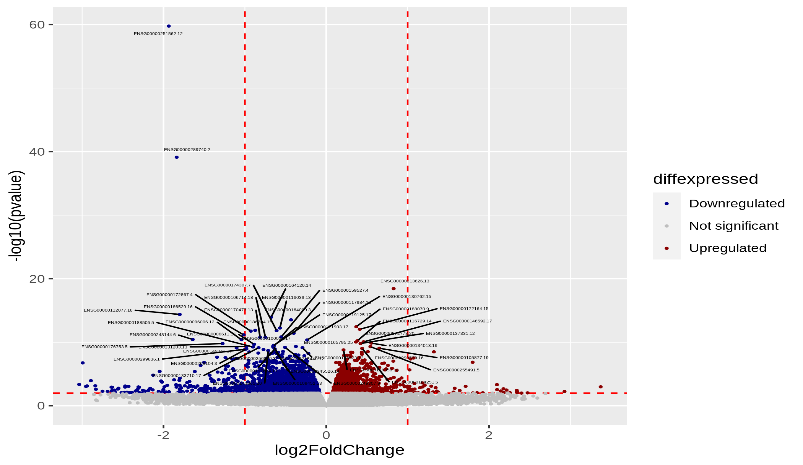

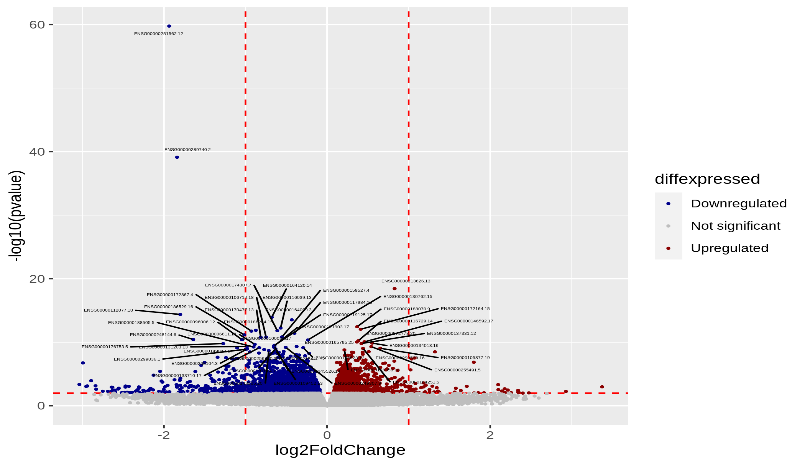

ASO1 vs ScASO

ASO2 vs ScASO

**A**

**B**

**C**

**Supplementary Figure 6: Differential gene expression upon *MALAT1* depletion in the primary tumor from PDO-X models**

(A) Volcano plots comparing differentially expressed genes (DEG) in DS115T PDO-X tumors upon *in vivo* *MALAT1* KD for each ASO compared to Scrambled ASO (ScASO). Red dots: upregulated genes upon *MALAT1* KD, blue dots: downregulated genes, grey dots: not significant. ASO1 and ASO2 – *MALAT1* targeting ASOs. (B) *MALAT1* expression and DEGs for NH048T PDO-X tumors. (C) *MALAT1* expression and DEGs for NH85TSc PDO-X tumors. Red arrow: *MALAT1* gene.

**Figure S7**

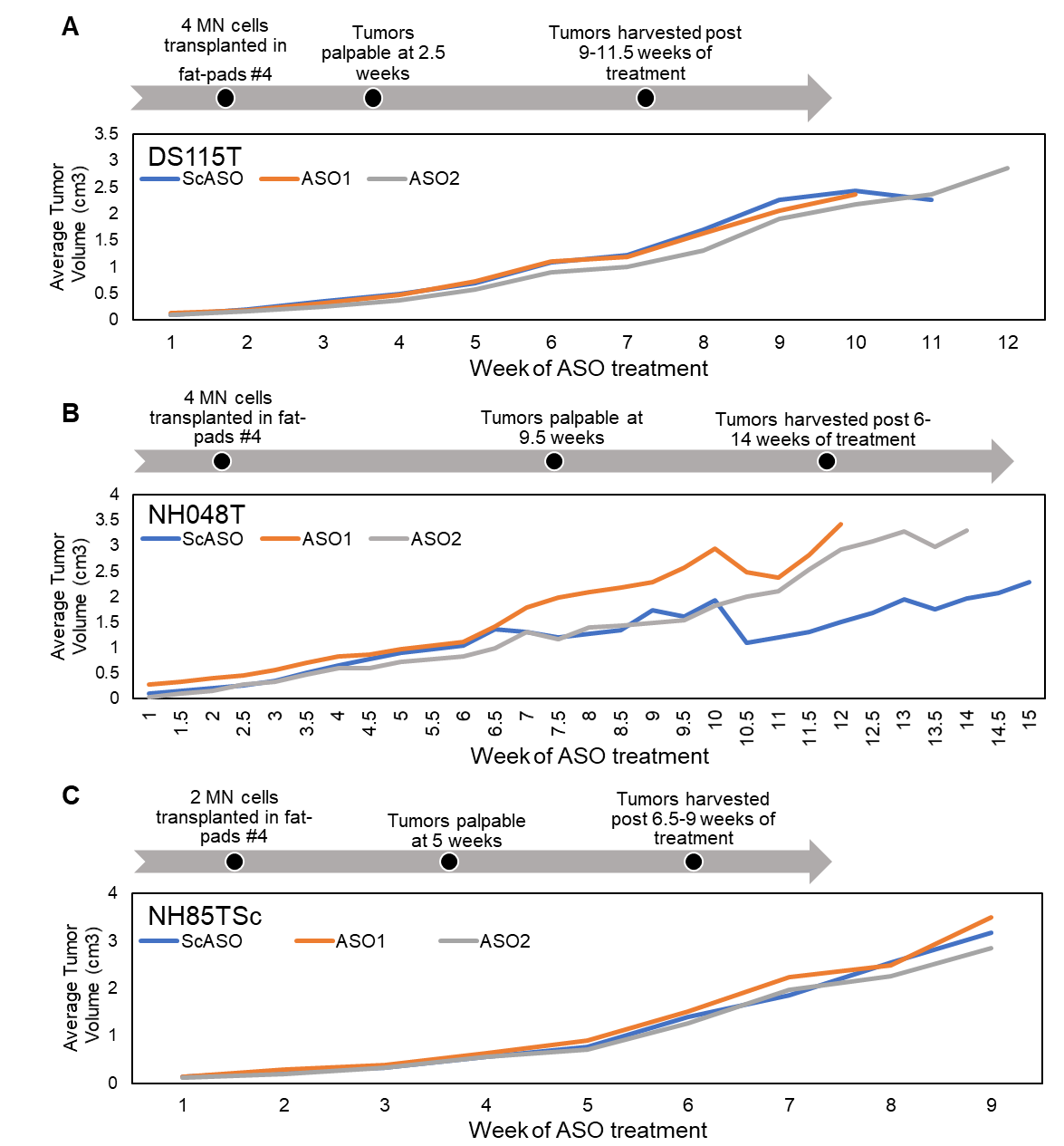

**Supplementary Figure 7: Treatment timeline and corresponding tumor volume over time upon systemic ASO (human-specific) treatment targeting *MALAT1* in the three PDO-X models.**

(A) Treatment timeline and corresponding palpable tumor volume change in the mammary fat-pad tumors of DS115T PDO-Xs upon systemic treatment with either a ScASO control or a *MALAT1*-targeting ASO (ASO1/ASO2). (B) Data representing treatment timeline and corresponding tumor volume change over time for NH048T PDO-Xs. (C) Data representing tumor volume change over time for NH85TSc PDO-Xs. Each curve represents an average tumor volume of 12-16 tumors (n=6-8 mice/group).

**Figure S8**

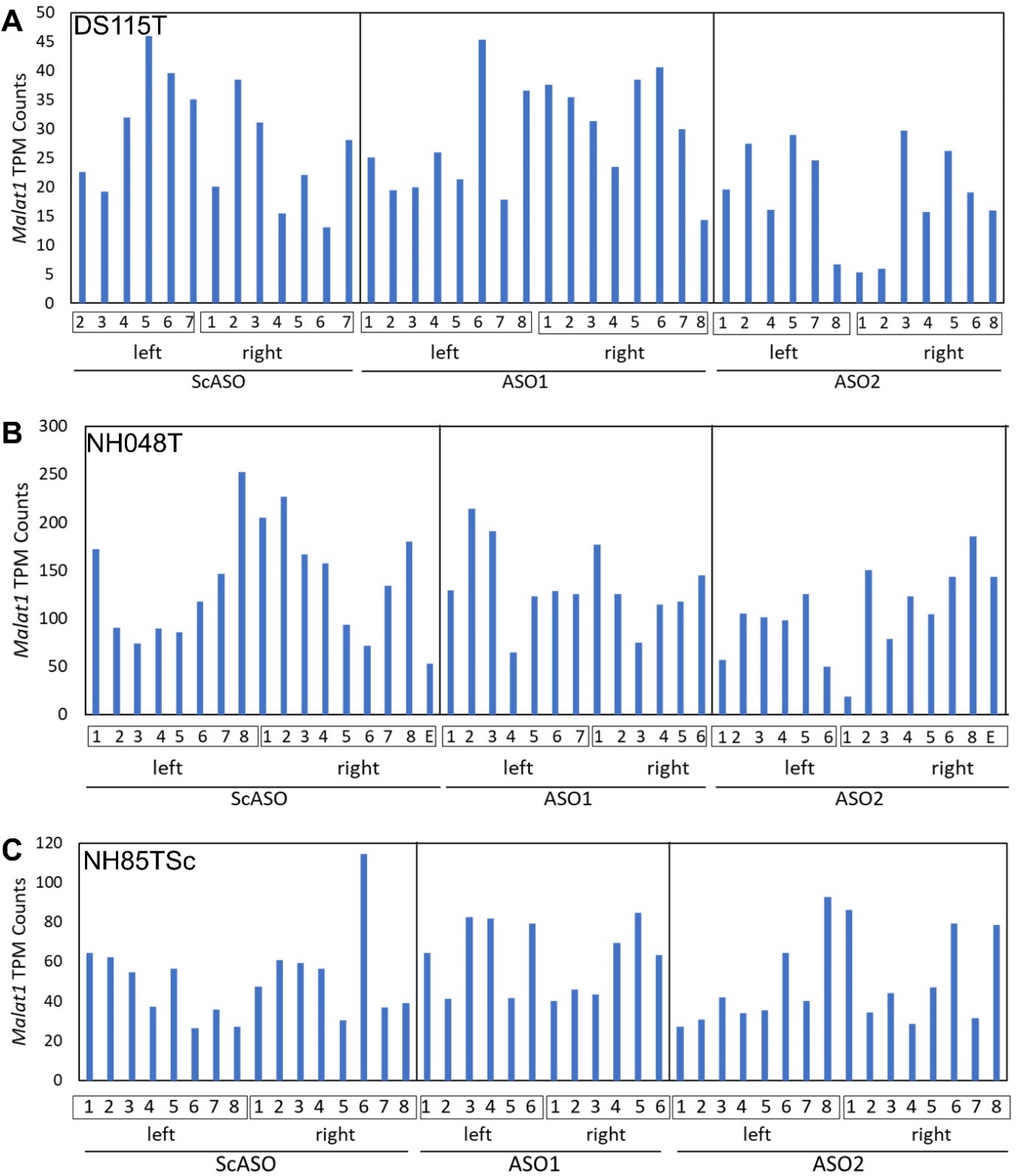

**Supplementary Figure 8:** ***Malat1* (mouse) transcript counts as measured via bulk RNA-seq in PDO-X-derived tumors.**

*Malat1* transcripts per million (TPM) counts for individual PDO-X-derived tumors for (A) DS115T, (B) NH048T and (C) NH85TSc PDO-Xs treated with either ScASO/ ASO1/ ASO2. Each number corresponds to animal # and left/right represent the tumor fat-pad location. E - additional separate tumor at one of the locations. ScASO – Scramble ASO control, ASO1 and ASO2 – *MALAT1*-targeting ASOs (human-specific).

**Figure S9**

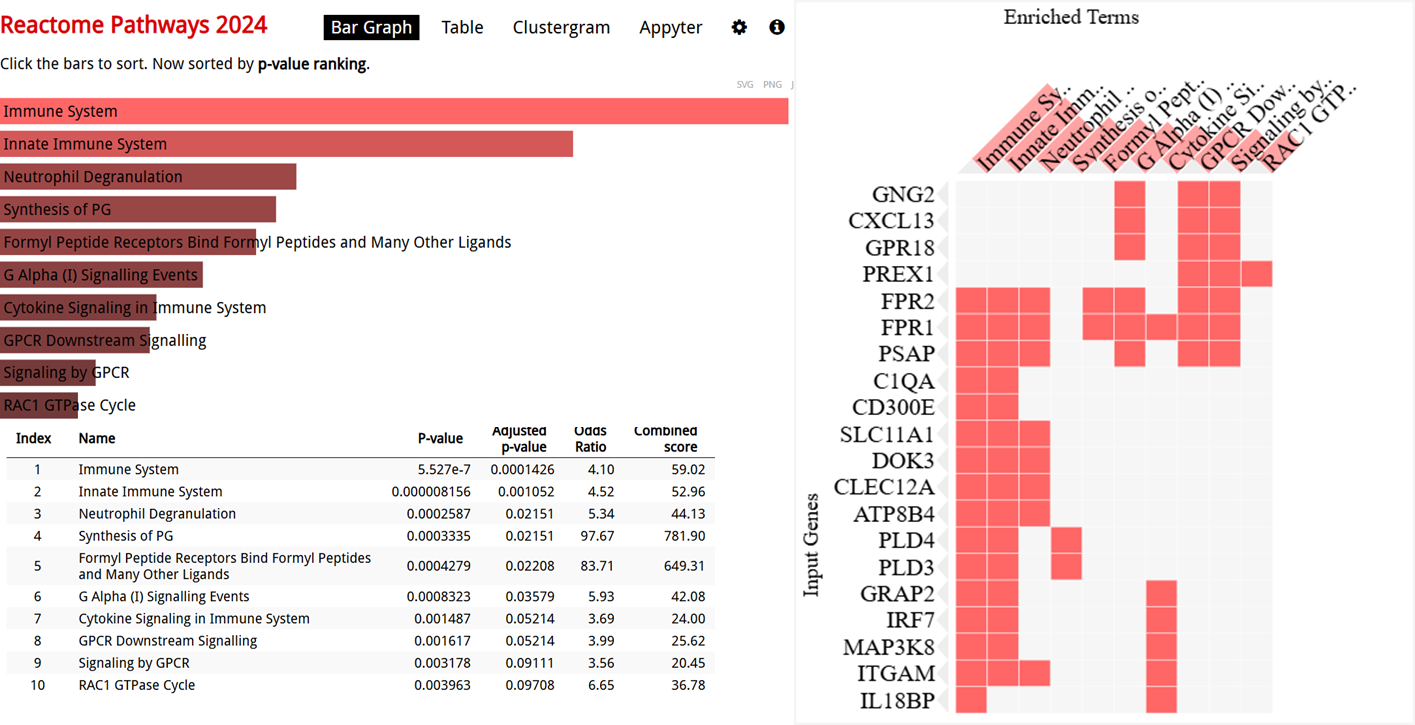

**Supplementary Figure 9: Pathway changes associated with *MALAT1*-driven common differential gene expression in the stromal compartment of three independent PDO-X tumor experiments.**

Top 10 enriched biological pathways corresponding to the set of commonly regulated stromal (mouse) genes upon *MALAT1* depletion across three independent (DS115T, NH048T and NH85TSc) PDO-X tumor studies, as observed in the Reactome 2024 database via ENRICHR.
